# Distinct replay signatures for prospective decision-making and memory preservation

**DOI:** 10.1101/2021.11.08.467745

**Authors:** G. Elliott Wimmer, Yunzhe Liu, Daniel C. McNamee, Raymond J. Dolan

## Abstract

Theories of neural replay propose that it supports a range of functions, most prominently planning and memory consolidation. Here, we test the hypothesis that distinct signatures of replay in the same task are related to model-based decisionmaking (‘planning’) and memory preservation. We designed a reward learning task wherein participants utilized structure knowledge for model-based evaluation, while at the same time had to maintain knowledge of two independent and randomly alternating task environments. Using magnetoencephalography (MEG) and multivariate analysis, we first identified temporally compressed sequential reactivation, or replay, both prior to choice and following reward feedback. Before choice, prospective replay strength was enhanced for the current task-relevant environment when a model-based planning strategy was beneficial. Following reward receipt, and consistent with a memory preservation role, replay for the alternative distal task environment was enhanced as a function of decreasing recency of experience with that environment. Critically, these planning and memory preservation relationships were selective to pre-choice and post-feedback periods. Our results provide new support for key theoretical proposals regarding the functional role of replay and demonstrate that the relative strength of planning and memory-related signals are modulated by on-going computational and task demands.

**Significance statement:** The sequential neural reactivation of prior experience, known as replay, is considered to be an important mechanism for both future planning and preserving memories of the past. Whether, and how, replay supports both of these functions remains unknown. Here, in humans, we found that prior to a choice, rapid replay of potential future paths was enhanced when planning was more beneficial. By contrast, after choice feedback, when no future actions are imminent, we found evidence for a memory preservation signal evident in enhanced replay of paths that had been visited less in the recent past. The results demonstrate that distinct replay signatures, expressed at different times, relate to two dissociable cognitive functions.

## Introduction

Humans have a remarkable ability to process information that extends beyond the immediately perceptible, including simulation of prospective plans and retrieval of past memories. It has been hypothesized that hippocampal replay contributes to both these abilities (1–5). In rodents, replay is strongly linked to the hippocampus, where cells encoding distinct locations reactivate in a coordinated sequential manner, recapitulating past or simulating potential future trajectories (1–6). A similar phenomenon of sequential reactivation has been identified in humans using decoding techniques in conjunction with high temporal resolution MEG data (7–14).

Prominent theories of neural replay propose that it is important for planning future actions (15–18) in addition to supporting memory preservation (19–25). One hypothesis is that task demands, operationalized as temporal proximity to action versus feedback, determine the contribution of replay to planning and memory, respectively (1, 26). However, the contribution of awake on-line replay to these two functions has largely been addressed in the context of separate experiments (1, 4, 6, 7, 26–33). Here we directly address the contribution of replay to both these roles within a single task context.

Replay of trajectories leading to a goal has been proposed to underpin decision making that exploits structure knowledge of an environment, referred to as model-based decision-making (15, 16, 34). A number of rodent studies indicate a link between hippocampal neural sequences and subsequent path choice selection, consistent with a role for replay in planning (6, 31, 35–37). However, an inconsistency in such findings raises the possibility that any relationship between replay and subsequent choice might differ across evaluation strategies and reward environments (6, 13, 31, 33, 35, 36, 38, 39). Critically, and regardless of any relationship to choice identity, brain lesion studies highlight a necessary role for the hippocampus in model-based behavior (40, 41). This suggests that hippocampal replay may be enhanced when model-based decisionmaking (planning) is beneficial. Thus far, however, there is no clear evidence linking demands for model-based control and neural replay preceding choice (9–11, 13).

Beyond planning, replay is considered critical for memory preservation, where replay of previous experiences might serve to strengthen memory and prevent interference from newer experiences (‘catastrophic forgetting’) (19, 20, 22, 42, 43). Studies that have disrupted hippocampal activity support the idea that offline place cell reactivation is critical for learning, memory consolidation, or both (44–47). It has been conjectured that human rest-period replay subserves a similar function (8, 12, 48, 49). Studies of replay in rodents navigating a single environmental context provide initial, but inconclusive, evidence for a link between recent experience with an environment and replay (28, 39).

Here, we address a role for replay in both planning and memory preservation in a context where participants needed to retain a memory of a ‘distal’ environment while at the same time learning within a local one. To do this we adapted a reward learning task originally designed to study model-based decision-making (41, 50–52), where distinct start states converge upon shared paths. Critically, to study memory, we included two independent randomly alternating environments. Both the early convergence on shared paths and the alternation of environments strongly favors online planning, and these features distinguish the current paradigm from a recent related report (11). Using recently-developed MEG analytic methods (7–9, 11–13, 53) we first identify sequential neural reactivation and then ask whether replay strength varies as a function of task demands and recent experience. We hypothesized replay would be boosted during prechoice path planning when model-based decision making was more beneficial (15–17, 40, 41, 50). By contrast, following receipt of choice feedback, we hypothesized replay for an alternative environment would relate to the infrequency of recent experience, consistent with a role in supporting memory preservation (1).

## Results

### Behavior

Participants navigated two separate, independent environments (‘worlds’) in order to earn reward points (**Fig. 1a**). Each world contained two path options, where a top-level shape led deterministically to one of the sequential paths comprising 3 unique stimuli.

**Figure 1.**
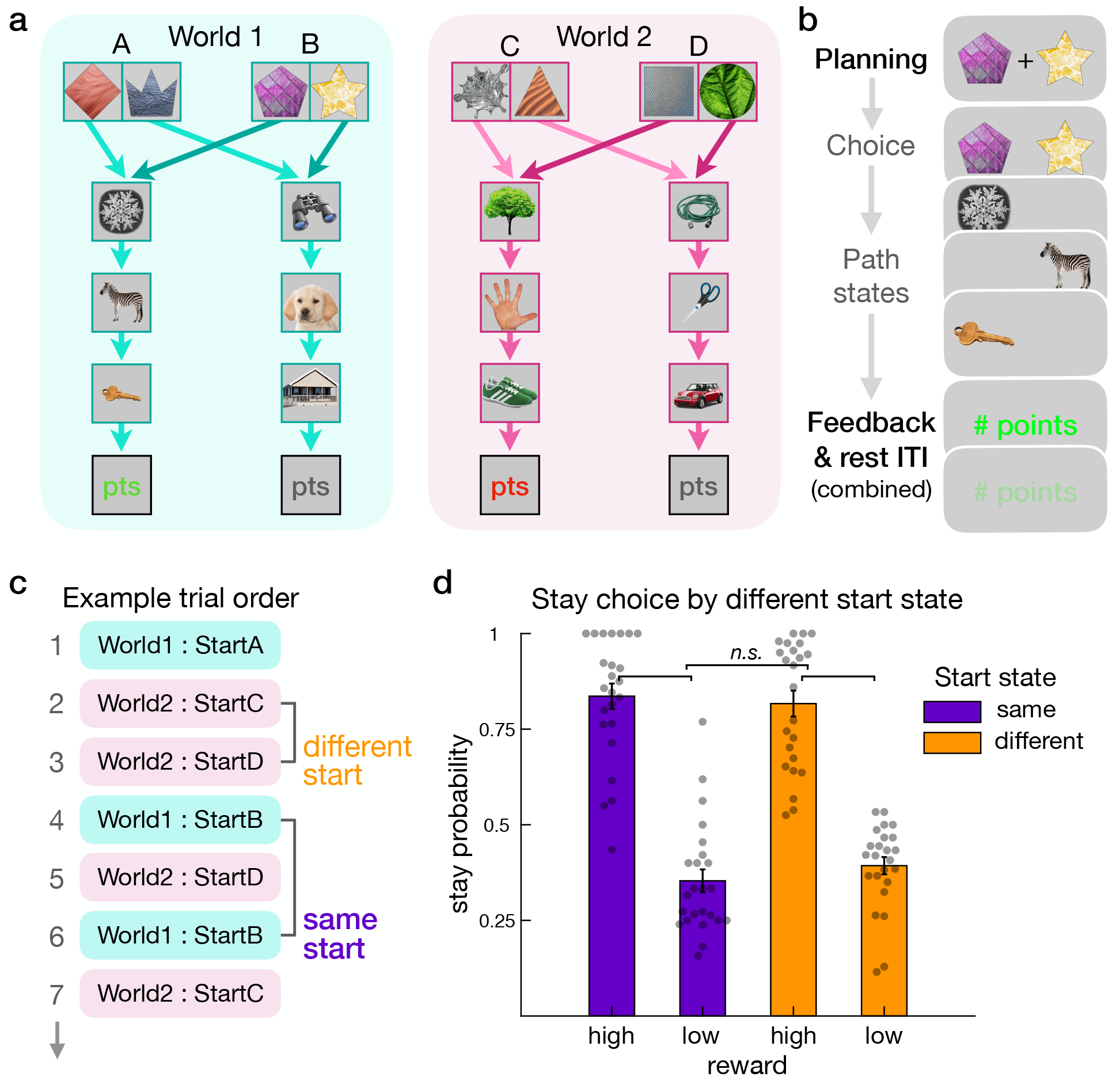
Two-environment reward learning task and generalization behavior. (**a**) Task schematic showing two alternative ‘worlds’ and their two equivalent start states. Trials in each world start at one of two equivalent shape pair options; to illustrate these connections, arrows from different states differ in color saturation. The shape options then lead deterministically to the same paths and reward outcomes (0-9 points (pts)). To learn this general structure participants engaged in a pre-scanning training session. For the MEG scanning session, participants then learned two worlds populated with new images. Participants’ memory for the path sequences was at asymptote after an initial no-reward exploration period. (**b**) Key trial periods in the reward learning phase. Replay was measured prior to choice in a time window we subsequently refer to as the ‘planning period’. After the disappearance of a central cross, participants entered their response. Participants then sequentially viewed the state images corresponding to the chosen path. Finally, participants received reward feedback (0-9 points), the amount of which drifted across trials. Replay was again measured in the post-feedback time window. For interpretation of subsequent results, in this example World 1 is the ‘current world’ while World 2 is the non-presented ‘other world’. (See also **Fig. S1**.) (**c**) Example trial sequence, highlighting two cases where a trial either has a different start state or the same start state as the previous trial in the same world. (**d**) Illustration of the dependence of repeated choices (stay) on previous rewards, conditional on whether the start state in the current world was the same as in the previous trial or not. The plot depicts the probability of a stay choice (when participants repeated a previous path selection in a given world) following above-average (high) versus below-average (low) reward. This difference was equivalent for same (purple) versus different (orange) starting states, consistent with behavior being model-based (‘n.s.’ represents non-significant effects in a regression model using continuous reward data). For display purposes, graded point feedback was binarized into high and low and trials with nearmean feedback were excluded; alternative procedures yield the same qualitative results. Grey dots represent individual participants. (See also **Fig. S2**.)

Knowledge of these paths was tested after each scanning block. In a given world, each path led to a separate stream of reward feedback which drifted over time (**Fig. S2a**). Importantly, within a given world, each trial could start in one of two equivalent start states leading to the two paths. For an agent to perform well in the task, outcomes should have an equivalent influence on subsequent choices regardless of whether an agent starts in the same start state or the alternative, equivalent, start state. This design feature, combined with a sufficient drift in reward across trials (51), allows us to characterize how well participants use structure knowledge to guide model-based behavior (**Fig. 1c**) (50, 51, 54, 55).

By way of example, imagine an agent is faced with a choice in world 1, start state A (**Fig. 1a**). The agent selects the diamond over the crown and receives an unexpected high reward. If next faced with a choice in start state B, a model-based agent will generalize this experience to promote the subsequent choice of the magenta pentagon to reach the same just-rewarded path. In contrast, a model-free agent does not have access to this recent relevant experience. Thus, only a model-based agent can exploit structure knowledge to allow generalization of reward feedback across equivalent start states. Such behaviour has been proposed to involve looking ahead to values associated with terminal states, possibly using prospective neural reactivation or replay (50). Thus, this design allows us to identify behavior reflective of model-based versus model-free learning (50, 51, 54, 55), analogous to variants of the paradigm that use probabilistic state transitions (40, 41, 52).

One feature of our deterministic task variant is that trials are divided between those where model-based behavior is beneficial (different start state) versus neutral (same start state) (50), allowing us to test for an association between neural replay and the benefits accruing from model-based behavior. Furthermore, a deterministic transition structure and an absence of branching paths increases our ability to detect evidence of sequential neural reactivation (11). With respect to planning, the fact that our design includes multiple worlds decreases the predictability of an upcoming trial, thus promoting deployment of planning-related processes at choice as opposed to after outcome feedback (11, 56). Critically, including multiple worlds also allows us to examine replay signatures of memory preservation for more distal (non-local) experiences.

After an initial scanned exploration period without reward feedback, participants performed the primary reward learning task. To ascertain the degree to which behavior was guided by model-based and model-free learning we used a regression approach in combination with a set of reinforcement learning models. Our regression analyses quantify a model-based influence on behavior by testing for an effect of generalization: whether a prior reward has a different effect on choice when starting in the same start state versus a different state than previous experience (50, 51). A model-based controller acts to generalize reward feedback across equivalent starting states, potentially using structure knowledge to look ahead and evaluate expected terminal rewards. Alternatively, updating the equivalent start state can occur after feedback (prior to choice), a form of non-local learning (11). In our analysis, if the effect of previous reward on choice is similar when starting in a different versus same state, then this indicates that participants’ behaviour reflects an exploitation of structure knowledge for a task environment (50, 52). Qualitatively, we observed that receipt of high versus low reward enhanced the likelihood of choice repetition (**Fig. 1d**) while a high degree of model-based behavior was evident in equivalent stay probabilities for choices starting at the same versus a different state (**Fig. 1d**).

Using logistic regression we quantified the effect of reward and start state on choice (51), finding a robust overall effect of previous reward on stay choices (β (regression coefficient) = 0.536 [0.424 0.647]; z = 9.434, p <0.0001). Importantly, there was an equivalent influence of previous reward on stay choices in the same versus a different start state, consistent with the absence of a significant model-free contribution (interaction between reward and same start state, β = 0.0313 [-0.0365 0.0991]; z = 0.905, p = 0.365). As a non-significant effect does not provide evidence in support of the null hypothesis, we employed a two-one-sided test (TOST) equivalence procedure to enable us to reject the presence of a medium- or larger-sized effect (57). Indeed, based on this we can reject a medium- or larger-sized model-free interaction effect (TOST equivalence test p = 0.027). While reward-guided choice behaviour was unaffected by start state changes, we found that reaction times were overall slower for choices on start state change trials (different versus same start state trials β = 0.009 [0.0017 0.0163]; z = 2.423, p = 0.0155), even though the delayed choice limits reaction time variance. In a complementary regression approach, testing the effect of previous reward on the identity of option selection (**Supp. Results**) (50), we again found no interaction between reward and start state, consistent with a model-based learning signature (50, 52). Overall, this allows us to infer that behavior is guided by reward to the same degree irrespective of generalization demands, likely reflecting participants’ behavior being strongly model-based.

Next, we compared fully model-based and model-free reinforcement learning models to a hybrid model commonly used to assess the relative strength of model-based and model-free learning (51, 52). The hybrid model includes a weighting parameter *w* which controls the degree of model-based influence on choice. Overall, behavior was best explained by a model-based controller, which outperformed the hybrid model, while the fit of the model-free controller was poor (**Table S1**-**S2**). While the hybrid model exhibited a numerical benefit in raw fit, when penalized for model complexity the pure modelbased controller provided an equivalent (measured via AIC) or better fit (measured via BIC; **Table S2**). At the individual participant level, the model-based controller provided a better fit for more than 83% of participants (using either AIC or BIC). Finally, given that good performance requires model-based generalization throughout the task, as expected we found no evidence for a change in model-based behavior over time (**Supp. Results**). The high degree of model-based behavior provided a robust context for us to next examine the associated neural processes.

We also obtained memory tests for sequential path stimuli to confirm that participants learned the path structure. Our task was designed to achieve high memory performance by the start of reward learning, as this would serve to both boost model-based behaviour as well as our ability to detect sequential reactivation. Memory performance, already at a high level at the end of the incentivized structure learning phase (second half, 87.5% [78.2 96.8]), was maintained at this level across the reward learning phase (mean 95.0% [91.3 98.7, with all blocks above 92%; effect of block, p > 0.64).

### Sequenceness identification

In our neural analyses, we first established reliable decoding from neural patterns evoked by the unique stimuli that indexed individual path states (**Fig. 1a**). A classifier trained during a pre-task localizer showed successful discrimination of all path stimuli, with peak decoding evident from 140-210 ms post stimulus onset. Based on this, and to maintain consistency with our prior studies, we selected a post-stimulus 200 ms time point for subsequent replay analyses (7–9, 11). The trained classifier generalized from the localizer to the actual presentation of path objects during reward learning, showing significant across-task classification (t_(23)_ = 7.361, p < 10^-7^; **Fig. 2** and **Fig. S4**). We also found evidence for significant reactivation of stimuli both during planning and after feedback (as compared to reactivation derived from permuted classifiers; p-values < 0.01; see **Methods**).

**Figure 2.**
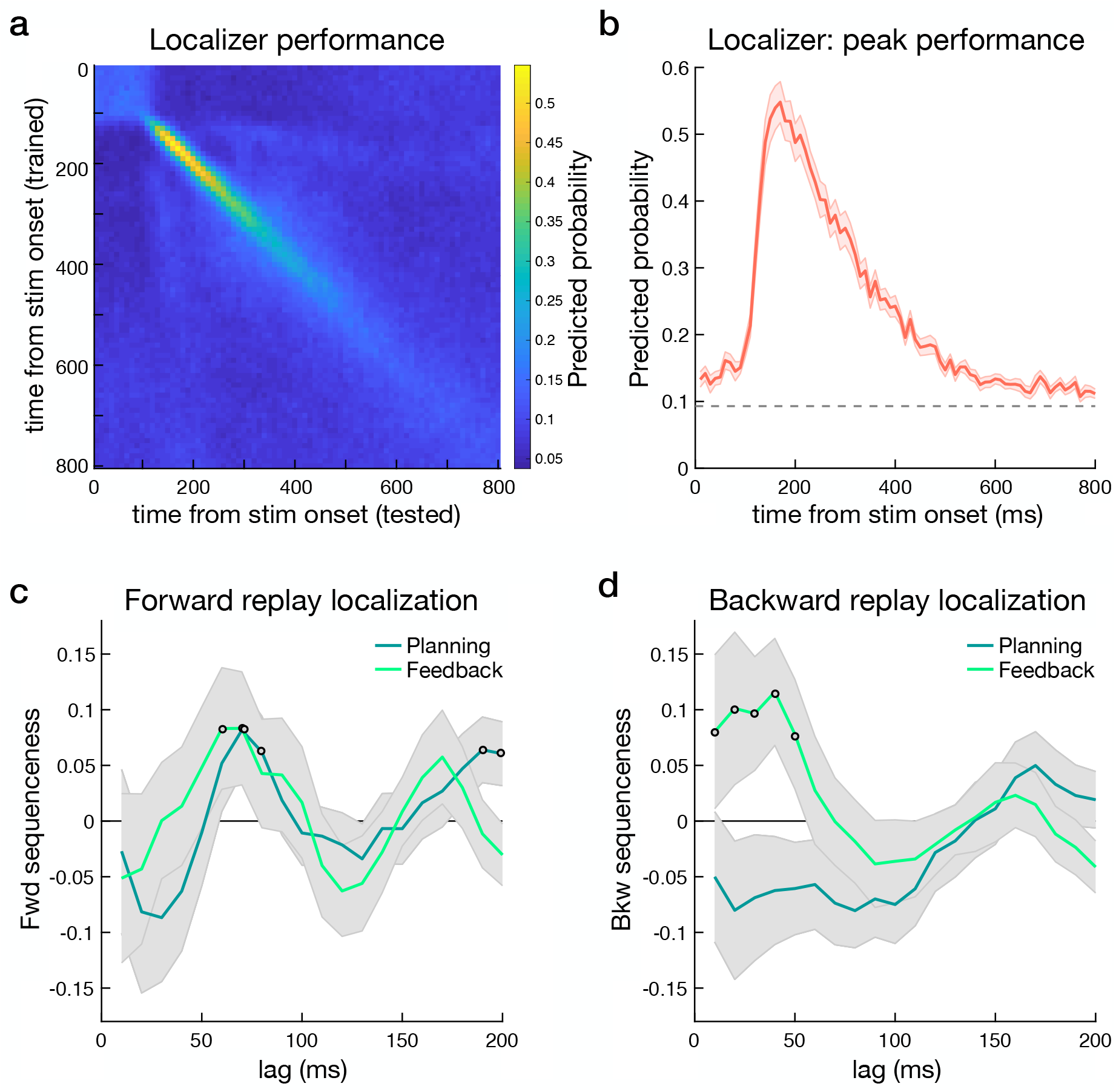
Training of state localizer and sequenceness time lag identification. See also **Fig. S3**. (**a**) Classifier performance for path state stimuli presented during a pretask localizer phase, training and testing at all time points. This revealed good discrimination between the 12 path stimuli used in the learning task. Color bar indicates predicted probability. Note that start state shape stimuli were not included in the pretask localizer and are not included in sequenceness analyses. (**b**) Peak classifier performance from 140-210 ms after stimulus onset in the localizer phase (depicting the diagonal extracted from panel a). (**c**) Forward sequenceness for all learned paths during planning and feedback periods was evident at a common state-to-state lag of 70 ms in both trial periods. Open dots indicate time points exceeding a permutation significance threshold, which differs for the two periods. (**d**) Backward sequenceness for all learned paths during planning and feedback periods was evident at state-to-state lags that spanned 10-50 ms in feedback period alone. Note that the x-axis in the sequenceness panels indicates the lag between reactivations, derived as a summary measure across seconds; the axis does not represent time within a trial period. Open dots indicate time points exceeding a permutation significance threshold, which differs for the two periods. Shaded error margins represent SEM. See **Fig. S6** for example sequenceness events and **Fig. S5** for extended time lags.

We next used the trained classifiers to seek evidence for time-compressed sequential reactivation of path elements. First, we applied the classifiers to reward learning task MEG data to derive measures of state reactivation in each trial separately for each state and at each time point, focusing on pre-choice planning and post-feedback rest periods (**Fig. 1b**). Next, we tested for time lagged cross-correlations between state reactivations within these periods, yielding a measure of ‘sequenceness’ in both forward and backward temporal directions at each lag (7, 8, 53) (**Fig. S3**). We use the term sequenceness (or ‘sequence strength’) to refer to the prediction strength of state *j* to state *i* at some time lag, while we operationally refer to any reactivation of state sequences here as replay. This sequence detection method, validated in previous work, quantifies the average predictivity of state *j* to state *i* within a period, reflecting both the frequency and fidelity of replay events (7, 8, 11, 12, 53) (**Fig. S3**).

Our a priori hypothesis is that signatures of replay are differentially modulated by task environment and recency of experience with an environment; the presence of forward or backward replay during planning and feedback (e.g. 2, 9) is a necessary precondition for testing this hypothesis. An initial temporal lag localization step independently identified lags of interest for subsequent examination of links between replay and behaviour. To increase power for localization, sequences included all possible paths, the (to-be) chosen and (to-be) nonchosen paths in the current trial, as well the two paths for the ‘other world’. We identified sequenceness time lags of interest by comparing evidence across lags for all valid sequences, using a significance threshold determined by a permutation of stimulus assignment to paths (following previous work; 8, 9, 11, 53). These analyses revealed that during planning there was significant forward sequenceness, with a state-to-state time lag of 70 and 80 ms and also of 190-200ms. For the shorter lags, this indicates that, on average across participants, a given state was followed by reactivation of an adjoining state within the same path at a delay of 70 to 80 ms (**Fig. 2c** and **Fig. S5**). We found no significant evidence for backward sequenceness during planning.

We then examined replay following outcome feedback, a period when the displayed reward points faded from the screen toward a brief inter-trial interval rest (**Fig. 1b**; **Fig. S1a**). This corresponds to a time when replay has previously been identified in rodents (e.g. 30). Here we identified significant sequenceness with a peak state-to-state time lag at 40 ms in a backward direction, and 60 to 70 ms in the forward direction (**Fig. 2c-d** and **Fig. S5**). To focus on a period with less cognitive demands arising from actual feedback processing and value updating, with similarities to a procedural step in a related rodent study (26), our analysis was focused on the latter 3.5 s of the 5 s post-feedback period; note, however, qualitatively similar results were found when using the full feedback period. In general, our finding of forward and backward replay events, intermixed across seconds, echoes results reported in rodent studies (38, 58) as well as in a recent human study (11).

Based on these initial replay temporal lag localization analyses, we focused our primary analyses on forward sequenceness with a 70 ms lag between states identified in both the planning and feedback periods. At feedback, we selected the peak lag of 40 ms from the above-threshold lags for backward sequenceness analyses, informed by our previous work (8, 9). To examine links between replay and task experience, we estimated sequenceness separately for path transitions in the current world and other world, for each period on each trial.

### Replay during planning and model-based generalization

We first tested a link between planning and relative sequence strength, leveraging the fact that in our task model-based generalization was more beneficial when a start state changes relative to when it remains the same (**Fig. 1d**) (50). A planning account predicts that when the start state changes the benefit of model-based generalization should be reflected in a boosting of replay relative to when the start state remains the same (**Fig 1c**). To increase power, our sequenceness analyses combined evidence for the two current world paths.

In line with predictions, sequenceness for current world paths was significantly stronger when the start state changed versus remained the same (i.e. when generalization was likely to be more beneficial; multilevel regression β (regression coefficient) = 0.1411 [0.0625 0.2197]; z = 3.519, p = 0.0004; **Fig. 3a**). There was no relationship between sequenceness for other world paths and generalization (β = −0.0201 [-0.0979 0.0577]; z = −0.506, p = 0.613; other world TOST equivalence test p = 0.012; current versus other difference z = 2.829, p = 0.0023, one-tailed). Analyses across different state-to-state lags indicated the presence of a selective effect of generalization centered on the independently selected lag of 70 ms (**Fig. 3b**). Further, the effect of generalization was stable across trials (positive interaction with trial, t = 1.055, p = 0.292; see **Supp. Results** and **Table S3** for additional control analyses). This relationship was also qualitatively similar for both the to-be chosen and the to-be non-chosen paths (p-values < 0.033). In follow-up control analyses, we also tested whether higher pre-choice replay was better explained by our planning account versus an alternative memory retrieval account. The planning account predicts a stepwise increase in replay on trials when generalization was more beneficial, whereas an alternative memory retrieval account predicts a graded increase in replay related to how far in the past a start state was last seen. We found that a planning account provided a better fit to the data (**Supp. Results**). Overall, the link between replay and model-based generalization rests on the detection of evidence for the same sequential neural representations being triggered by two different sets of choice cues, demonstrating a form of neural generalization underlying behavioural generalization.

**Figure 3.**
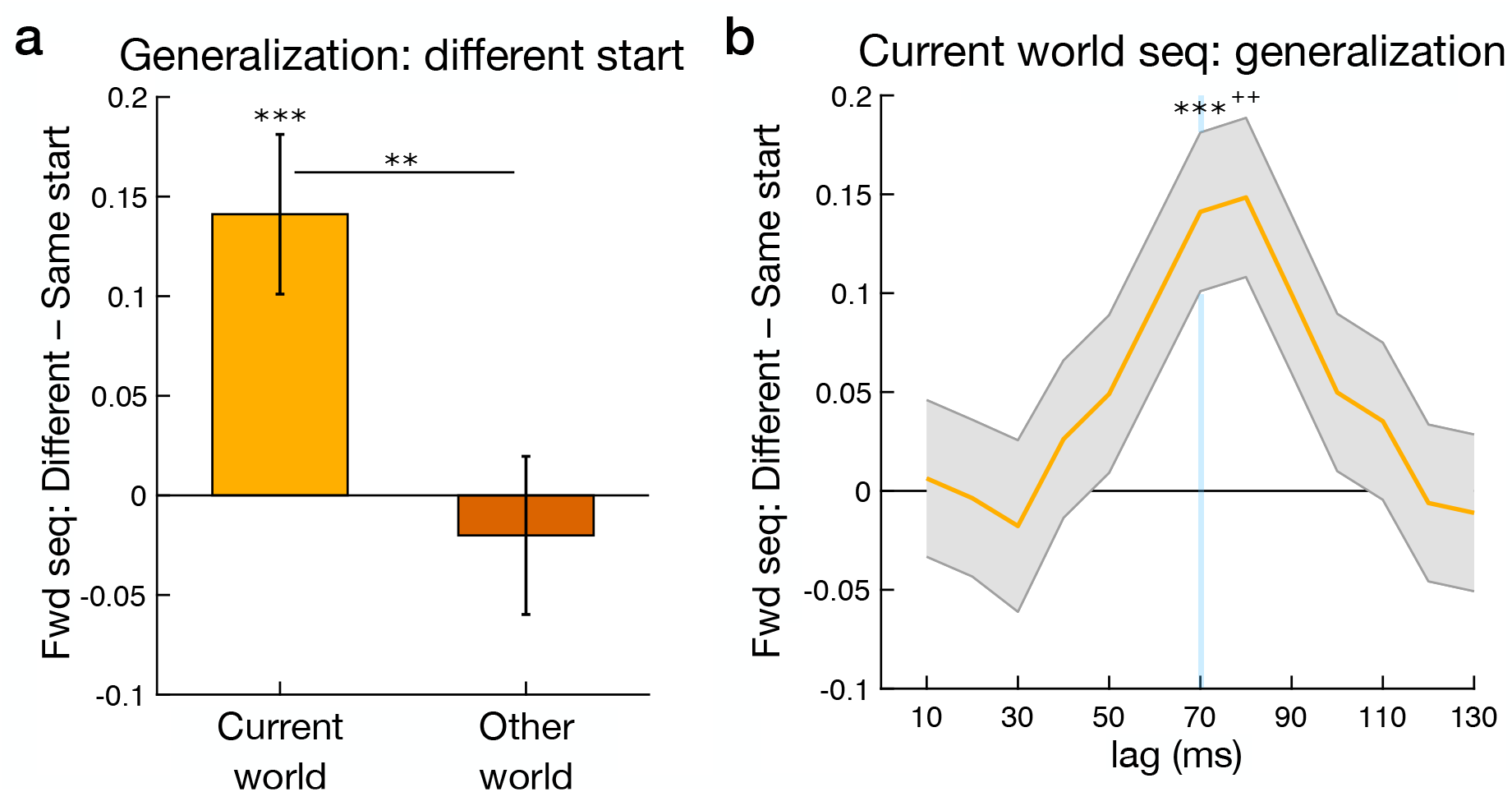
Planning period replay increases and the benefit of model-based generalization. (**a**) Stronger forward replay on trials where the start state is different from the previous trial, where there is a greater benefit in utilizing model-based knowledge (**b**) Timecourse of regression coefficients for variables of interest, showing effects at state-to-state lags from 10 to 130 ms. The light blue line highlights the 70 ms time lag of interest shown in panel a. Y-axes represent sequenceness regression coefficients for binary different versus same start state. See **Fig. S5** for extended time lags. Seq = sequenceness. ** p<0.01; *** p<0.001; + p<0.01, corrected for multiple comparisons.

We next examined whether planning-related replay was modulated by option value, in light of previous imaging and electrophysiology studies in humans reporting correlations between non-sequential hippocampal activity and value (e.g. 59, 60-62). If replay is involved in deriving value estimates, then we would not necessarily expect a modulation of replay by value, though it is possible that replay might be biased by option value when values are directly informed by recent experience.

Examining the relationship between replay and mean state value, the average model-predicted value across the two options, we found that current world replay strength significantly correlated with mean state value (β (regression coefficient) = 0.0479 [0.0130 0.0829]; t = 2.688, p = 0.0072). However, this relationship was only found on trials where there were no generalization demands (where the start state remained the same; same start trial value β = 0.1128 [0.0484 0.1771]; t = 3.440, p = 0.006; different start trial value β = 0.0110 [-0.0310 0.0530]; t = 0.513, p = 0.608; interaction β = 0.0510 [0.0140 0.0880]; t = 2.702, p = 0.0069). These results were also selective to current world forward replay during planning (**Supp. Results** and **Table S3**). Thus, replay positively related to option value in conditions where start state options were susceptible to direct reinforcement on the preceding trial, where we speculate value inference is less demanding.

### Backward replay prioritization at feedback and memory preservation

We next tested a prediction that replay during periods of low cognitive demand, specifically following reward feedback, relates to automatic memory maintenance processes, which we refer to as ‘preservation. Here we focused our analyses on backward replay with a 40 ms time lag, a signal selective to the feedback period (**Fig. 2d**). Behaviorally, we found consistently high levels of path memory during reward learning (see above) and this precluded examining direct links between feedback replay and memory variability. However, in the preceding brief structure learning phase when participants first experienced the sequential paths mean memory performance was lower (78.6% [68.0 89.3]), allowing us to test for a link with backward replay within the inter-trial interval (as no reward feedback was presented). Backward replay exhibited a numerical, but non-significant, increase across time (40ms lag, second half – first half trials; 0.101 [-0.034 0.237]; p = 0.136). Notably, increased backward replay from early to late trials in this initial phase correlated significantly with individual differences in memory performance during this phase (r = 0.409, p = 0.0470; **Fig. S5**). Although the number of trials here is much lower than the primary reward learning phase, this provides initial evidence consistent with a link between replay and memory.

After initial experience, memory preservation can be considered to be an automatic process driven in part by the infrequency, or rarity, of recent experience for a given environment (43). In the primary reward learning phase, we operationalized rarity as an exponentially weighted average of past exposures to each environment. We found that backward replay of other world paths was greater when they had been experienced less frequently over recent trials (rarity effect for other world paths, all trials β = 0.0513 [−0.0921 −0.0104]; t = 2.463, p = 0.0139; **Fig. 4a**). We found no relationship between current world replay and rarity (current β = −0.0101 [-0.0307 0.0509]; t = 0.486, p = 0.627; TOST p = 0.011; other versus current difference z = −2.076, p = 0.0190, one-tailed; **Fig. 4a**). Moreover, the relationship with rarity was stable across trials (interaction with trial, t = 0.487, p = 0.627; see **Supp. Results** and **Table S3** for additional control analyses). We also confirmed this experience-replay relationship in a basic model that makes no assumptions about learning, finding that other world replay was stronger when the other world had been experienced more than one trial ago versus when experienced on the previous trial (p = 0.0469).

**Figure 4.**
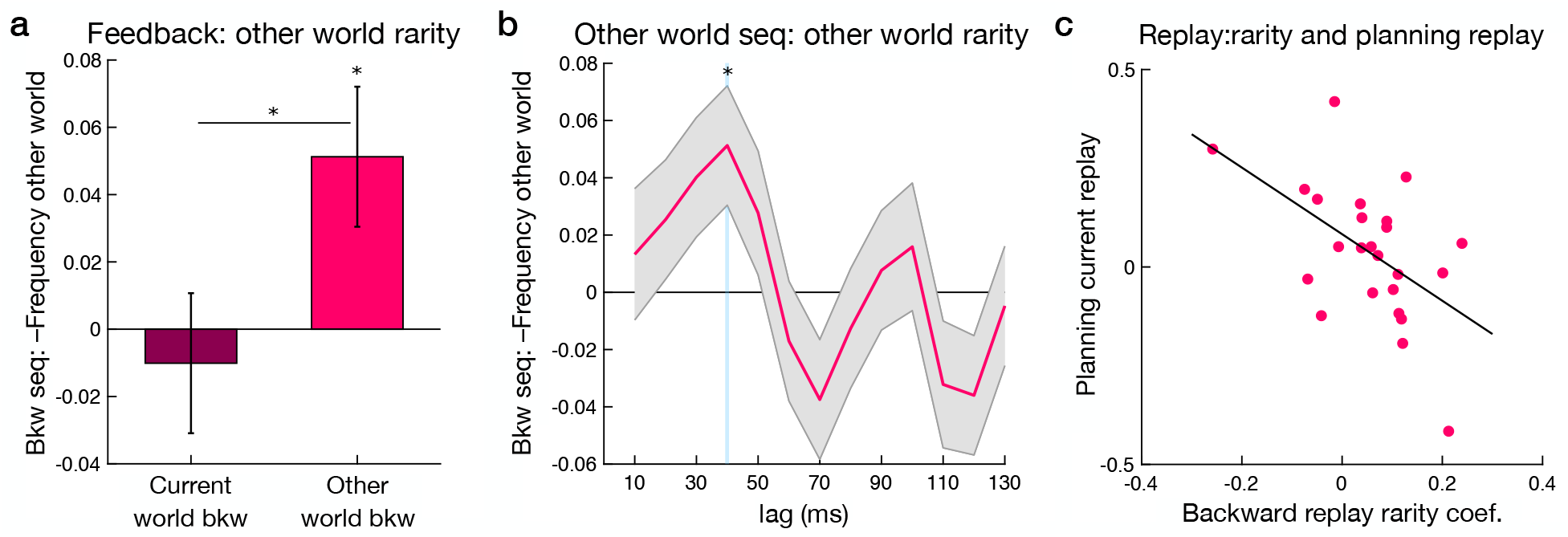
Feedback period backward replay increases with the rarity of recent other world experience. (**a**) Rarity (lower recent experience) of the other world correlated with greater backward replay of other world paths. (**b**) Timecourse of regression coefficients for the rarity effect of interest, showing effects at state-to-state lags from 10 to 130 ms. The light blue line highlights the 40 time lag of interest shown in panel a. Y-axes represent sequenceness regression coefficients for rarity of the other world. See **Fig. S5** for extended time lags. Seq = sequenceness; * p < 0.05. (**c**) Across-participant relationship between the replay-rarity effect and lower planning period forward replay (world change trials; p = 0.009).

To further explore this putative memory preservation signal, we also examined a link between the rare experience replay effect and planning forward replay. Stronger replay of rare experiences after feedback might be expected to decrease the need for planning replay. The planning replay signal was extracted from trials where the world changed from trial-to-trial alone, as this captures where any preceding feedback period ‘other world’ replay effects may relate to planning. Consistent with this we found an inverse relationship between the strength of the modulation of backward replay by rarity and planning replay across participants (current world forward replay, world change trials r = −0.521, p = 0.009; **Fig. 4**).

Next, we asked whether there was a link between this rare experience replay signature and choice behaviour. If feedback replay supports memories for more distant structure and value, we might expect the strength of this replay signal to positively influence choice in the other world. In an augmented reinforcement learning model, we tested this connection by allowing replay-related memory preservation to decrease choice uncertainty (or noisiness). The feedback replay measure was extracted from trials preceding a world change, as this is where preceding feedback period replay may relate to a following choice. The model included two additional softmax inverse temperature parameters that applied to world change trials with high versus low preceding feedback replay (**Methods**). A higher inverse temperature parameter can reflect lower uncertainty such that choices are more strongly guided by estimated prospective values. We found a significantly higher inverse temperature when choices were preceded by high replay versus low feedback replay (world change trials; high replay median = 19.40; low replay = 14.27; z = 2.171, p = 0.015, one-tailed, Wilcoxon signed rank test). Control analyses demonstrated that the replay effect on choice was selective to backward replay of the relevant world (**Supp. Results**). Further, while backward replay related to the experienced rarity of a world, we found no modulation of choice noisiness by experienced rarity itself, consistent with the internal variability of backward replay underlying the observed effect. Thus, supporting a potential memory preservation mechanism, we found 1) that backward replay was positively modulated by rarity of experience, 2) that a stronger rarity replay effect was linked to lower planning replay strength, and 3) that the strength of backward replay on a trial-to-trial basis decreases uncertainty in subsequent choices.

We then compared the task and experience links to replay that we identified during planning and after feedback to determine if these signals were distinct. We found no correlation between planning period forward replay and rarity of recent experience effect (current world 70 ms lag, p = 0.815), with the feedback period significantly stronger than the planning period effect (difference, z = 1.778, p = 0.0378, one-tailed). Conversely, we found no significant correlation between feedback period backward replay and the benefit of generalization (other world 40 ms lag, p = 0.119), while the planning period effect was significantly stronger than the feedback period (difference, z = 2.891, p = 0.002, one-tailed). Together, these planning and feedback comparisons represent a double dissociation, with feedback period replay being selective for an expected signature of memory preservation.

In additional control analyses, we found no relationship between backward replay and reward feedback or reward prediction error (**Supp. Results** and **Table S3**). Further, during planning, we identified significant forward replay with a 190 and 200 ms time lag (**Fig. 2c**), but found no correlation between this signal and any variables of interest (**Table S3**). At feedback, we also identified significant forward replay with a 70 ms time lag (**Fig. 2c**), but found no correlation between current world forward replay with feedback or other variables of interest (**Supp. Results** and **Table S3**). Finally, in exploratory analyses of a longer 160 ms lag replay signal identified recently (11), we found no relationship with variables of interest at feedback or during planning (**Supp. Results**).

### Replay onset beamforming and time-frequency analyses

To explore the spatial source of sequenceness events we conducted supplemental beamforming source localization analyses to test whether replay onset is associated with increased power in the hippocampus, as previously found (7, 8, 11, 12, 63). As the interpretation of sources of MEG signal is complex, especially for putative deep regions such as the MTL, we emphasize the supplemental nature of these analyses. Candidate replay onsets were identified by locating sequential reactivation events at time lags of interest, applying a stringent threshold to these events, and conducting broad-band beamforming analyses (as in 7).

At both planning and feedback periods, we identified power increases associated with replay onset in clusters extending from peaks in the visual cortex into the hippocampus (**Fig. 5**; planning forward replay peak MNI coordinates [x, y, z] 20, −81, −2; p < 0.05 whole-brain permutation-based cluster correction; feedback backward replay peak; −10, −91, −7; **Fig. S7**; **Table S4**). Using a hippocampal ROI analysis, during the planning period there was a significant power increase at replay onset evident in the right hippocampus (30, −36, −4; p < 0.05, ROI permutation-based cluster correction). Within the feedback period, we found bilateral hippocampal power increases (right 30, −21, −12 and 26, −36, 0; left −25, −16, −17; p < 0.05, corrected). Similar whole brain and hippocampal ROI results were found when separately looking at planning period current world replay onsets and feedback period other world replay onsets (**Table S4**). These results, along with recent related reports (7, 8, 11, 12), support an interpretation that sequential reactivation is tightly linked to enhanced hippocampal activity.

**Figure 5.**
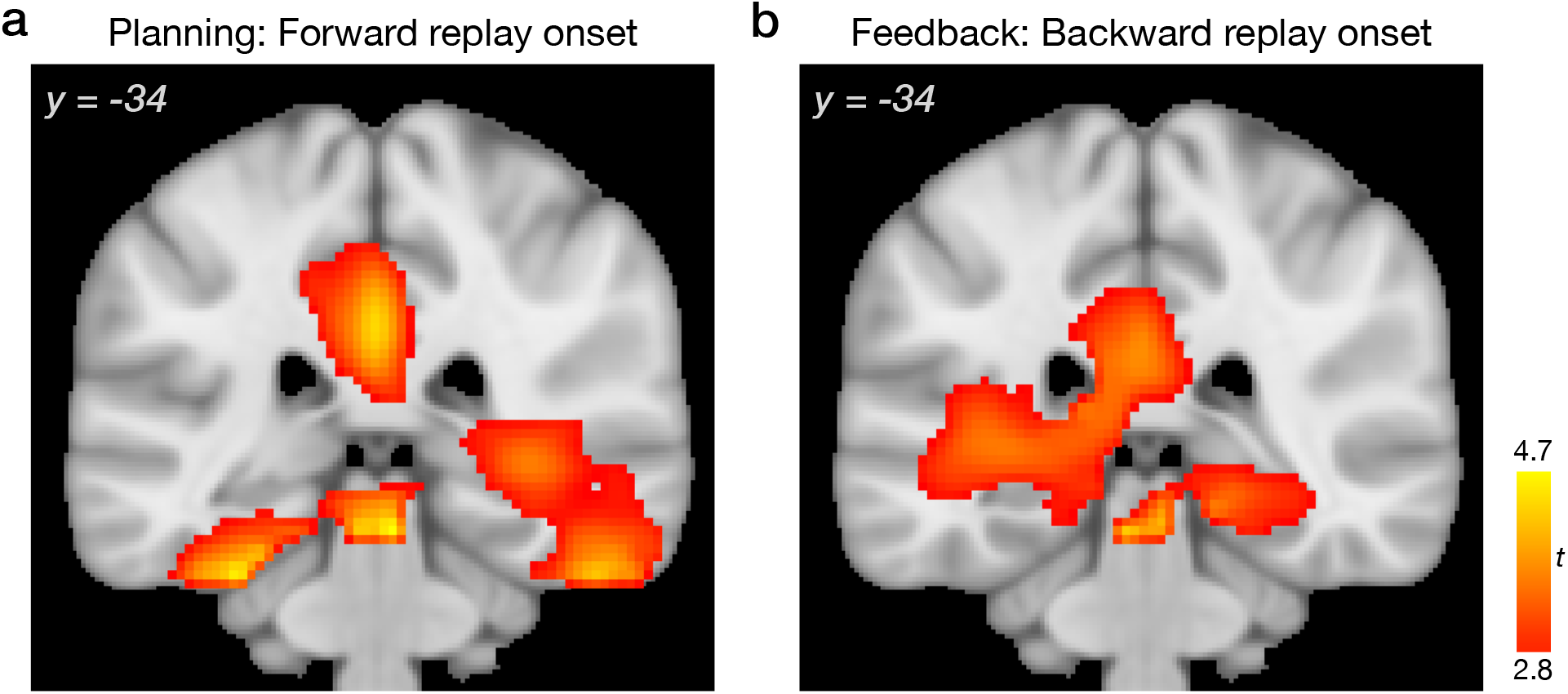
Exploratory replay onset beamforming analyses. (**a**) In the planning period, beamforming analyses revealed power increases associated with replay onset in the right MTL, including the hippocampus. (**b**) After reward feedback, power increases associated with replay onset were found in the bilateral MTL, including the hippocampus. See also **Fig. S7** and **Table S4**. The y coordinate refers to the Montreal Neurological Institute (MNI) atlas. For display, statistical maps were thresholded at P < 0.01 uncorrected; clusters significant at P < 0.05, whole-brain corrected using nonparametric permutation tests. For unthresholded statistical maps and results within the hippocampus ROI mask, see https://neurovault.org/collections/11163/.

In separate time-frequency analyses of power changes associated with replay onset events, we found that replay onset for both the planning and feedback periods related to increased power centered at 5-40 Hz (p < 0.05, permutation-based cluster-correction; **Fig. S8**). While we found no main effect of replay-related power increases in the ripple band (120-150 Hz) (8, 11, 12), at feedback, we found that individual differences in other world replay onset ripple power correlated with the strength of the modulation of backward replay by rarity (other world - current world; r=0.527, p=0.016 corrected for two comparisons; **Fig. S8**). A similar but non-significant relationship was found for the theta band (5-8Hz, r=0.449, p=0.0572, corrected).

### Generalized position representations across worlds

Finally, we asked whether learning led to changes in neural representations that reflected abstract, generalized, information about task structure (8, 12, 64). We predicted that neural representations of stimuli in the same path position (1, 2, and 3) would become more similar after learning (8, 12), an abstraction which might aid planning. We used Representational Similarity Analysis (RSA) to index representation changes from the pre-task localizer to reward learning path navigation (12, 65, 66).

When path stimuli were presented during learning, we found evidence for a significant representation of position information (**Fig. S9**; p< 0.05 FWE corrected). In particular, a position representation was evident for stimuli presented in different worlds (‘across-world’), consistent with position information generalizing across stimuli shown on separate trials. Thus, for example, stimuli occupying position 1 in world 1 showed greater similarity to stimuli in position 1 in world 2 than to stimuli in positions 2 or 3 in world 2. The effect was evident in an initial peak from 180–250ms (p < 0.0001 uncorrected), where across-world position information was present for all three individual positions when examined separately (p-values < 0.0012). Our design did not include a post-learning localizer, so these representations were necessarily assessed during reward learning. Because of this, it is possible that some position representation information during the learning task could arise from a preceding trial phase, or indeed from cognitive expectation effects. We note that while previous findings showed encoding of position information for stimuli interleaved during learning (8, 12), our results demonstrate that position information generalizes across structurally similar environments even where stimuli never overlapped.

We also investigated path identity representations. For decision-making, it can be helpful to differentiate between grouped stimuli so as to facilitate distinct reactivation during learning; alternatively, it may be helpful to increase similarity for grouped stimuli to allow for linking or chunking. Supporting a differentiation effect, path information emerged at 800ms after stimulus onset, as reflected in a significant decrease in similarity for stimuli in the same path (800-910ms, p < 0.05 FWE corrected; peak 820ms, p = p = 0.00038 uncorrected; **Fig. S9**). We did not find any relationship between position or path representation strength and behavioural or other neural measures; these null effects could partly be due to noise added by cognitive expectation effects during learning.

## Discussion

Two proposed roles for neural replay relate to prospective decision-making and memory preservation. We show that during decision-making, replay strength was related to the relative benefit of model-based, goal-directed, control of behavior. By contrast, after outcome feedback, replay of alternative environment paths positively related to the rarity in recent experience of a more distal environment. Furthermore, consistent with a putative role in memory preservation, stronger replay following feedback related to a subsequent decrease in behavioural choice uncertainty for the alternative environment. Thus, we find selective links between replay strength and the benefits of planning and the recency of experience, demonstrating distinct roles for replay within a single task.

By manipulating the benefit derived from model-based generalization of reward value across trials, we identified a relationship between planning replay and use of model-based inference. Building on previous imaging work which linked future state reactivation and individual variability in model-based behavior (13, 50) we show here, on a trial-by-trial basis, a boosting of replay when making a choice in a different, but functionally equivalent, state as on the preceding trial, potentially supporting value inference. Further, planning-related replay onsets were associated with power increases in the MTL consistent with a localization to the hippocampus. While previous lesion studies have demonstrated an overall role for the hippocampus in model-based behaviour (40, 41), our results suggest that this contribution occurs during model-based planning.

Model-based generalization can be accomplished using different strategies in addition to planning, such as updating ‘non-local’ options following feedback (11, 34, 56) or by forming overlapping representations through extensive experience. To illuminate planning, our design included two unique features. First, unlike a majority of learning experiments which repeat the same environment on each and every trial (e.g. 52, 56), we employed two separate environments, intermixed in an unpredictable manner across trials, thereby limiting the utility of planning for the next choice immediately after feedback. Furthermore, in our design, the two alternative start states in an environment converge upon shared paths at the very first step, potentially increasing the degree of inference required during planning to differentiate between trajectories. These two features, environment alternation and early path convergence, distinguish the current task from a recent report that focused on feedback linked replay signals (11), which revealed that generalization involved updating different non-local start states after feedback. Here, by contrast, we found no significant reward- or planning-related responses after feedback either for our replay signatures of interest, or for the signature identified in Liu et al. (11) (**Supp. Results**). Thus, our current results, and those of Liu et al. (11), both demonstrate a link between neural replay and model-based inference, but at different time periods. It is often the case that the decisions we face arise unpredictably, and our current results support the idea that replay is of benefit in such situations.

To successfully generalize reward feedback and perform well in our task, participants need a model-based strategy. However, similar to a design used in a recent related report (11), choices were only made at the first level, and thus evaluation of path steps was not strictly necessary for model-based behavior. Nevertheless, we found robust evidence for path replay, identifying a new link between replay and model-based decision-making (planning). Further, this relationship was specific to sequential replay and we found no effects related to the reactivation of individual states. We speculate that in many environments outside the lab, replay-assisted planning of trajectories is advantageous and that it may be a default strategy employed even when not strictly beneficial. Future experiments could usefully study these neural mechanisms in environments where sequential step-by-step choices are required, although a previous study in this domain did not identify connections to participants’ behavior (9). Similarly, to link sequential replay to value-guided choice, it will be important for future experiments to explore whether pre-choice replay events are linked to value estimates. Our task was not optimized for value decoding. However, in simpler contexts, replay has been linked to outcome representation during a post-learning rest period in humans (8), with related links reported between hippocampal activity and activity in the ventral striatum or ventral tegmental area in rodents (67, 68).

In contrast to planning, a rest period after outcome feedback entails minimal cognitive demands with respect to the current environment, rendering it likely that activity at this time-point might support preservation of weaker memories (1). Our results, selective to the feedback period, are consistent with this. Previous studies of hippocampal replay in rodents have suggested a link to less recent experiences (28, 39). However, in these studies experience was confounded with low value, and in one case, a shift in replay was observed even before a reduction in experience (39). By parametrically varying experience in two distinct environments and controlling for value, we provide a quantitative link between replay and infrequent experiences. Such a neural replay mechanism can act to reinforce memories that are at risk of becoming weaker, effectively serving as internally-generated interleaved training (22). While we found a link to memory preservation during ongoing behaviour, we speculate that such a preservation mechanism may also operate offline for memory consolidation.

Our experiment necessitated robust task structure knowledge and very high memory performance for the sequential paths. Behaviourally, previous research has shown that a strong understanding of task structure promotes model-based learning while, conversely, a poor understanding leads to idiosyncratic learning strategies (69). One potential limitation of high memory performance is that this precludes linking variability in memory performance and post-feedback replay. However, during reward learning, a memory preservation account does not necessarily predict a simple relationship between the two: if internal evidence of lower memory strength drives higher post-feedback replay, an effective replay mechanism may remediate any memory compromise before it can be observed. A feature of our design is that it allows us to identify a neural mechanism that may naturally assist in maintaining high memory levels for distant experiences. Further, our reinforcement learning models suggest a link between trial-to-trial strength in feedback replay and memory via lower noise in the following choice. In general, we speculate that replay-supported memory preservation supports adaptive behaviour in situations where memory is more variable, and these are avenues to explore in future experiments.

We suggest that there may be a trade-off between the two separate functions of planning and memory preservation that we identify. One idea is that the content and function of replay is modulated by task demands, which differ between planning versus resting after feedback (1). Consistent with this, it has been reported that replay content that is temporally proximal to active navigation is task-directed, while replay content during reward consumption is undirected, potentially related to preserving memory of the entire environment (26). Computationally, our results suggest the expression of replay may reflect an arbitration of resources as a function of a reliance on planning versus memory (70). In line with such a tradeoff, we found that across-participant strength of the memory replay effect related to a lower expression of planning period forward replay. Such arbitration between functions would also influence the degree to which replay supports updating of values, as in a recently proposed computational model of replay (11, 34). We did not formally manipulate task demands such as cognitive load during planning, but our results provide potential pointers for future targeted studies (71).

In conclusion, we provide evidence that prospective replay is enhanced when model-based behavior is beneficial, while replay consistent with memory preservation is observed when demands are low, consistent with distinct signatures for key proposed functions of neural replay (15–25, 49). While we dissociate these functions, both planning and memory functions are necessary for adaptive behavior, not least because a stable memory of the one’s environment aids successful decision making (1, 4, 72). By identifying replay signatures for planning and recency of experience, our results have relevance for targeting an understanding of common or separable disruptions to these functions in psychiatric disorders and disorders that impact on memory, such as Alzheimer’s disease (12, 73–77).

## Methods

Twenty-seven healthy volunteers participated in the experiment. Participants were recruited from the UCL Psychology and Language Sciences SONA database and from a group of volunteers who had participated in previous MEG studies. Of this participant group, the MEG session was not conducted in three participants: one due to scheduling conflicts, one due to technical problems with the MEG scanner, and one due to poor performance in the behavioral training session (see below). This resulted in the inclusion of data from 24 participants for behavioral and MEG analyses (14 female; mean age 23.8 years; range 18-34). Participants were required to meet the following criteria: age between 18-35, fluent English speaker, normal or corrected-to-normal vision, without current neurological or psychiatric disorders, no non-removable metal, and no participation in an MRI scan in the two days preceding the MEG session. The study was approved by the University College London Research Ethics Committee (Approval ID Number: 9929/002). All participants provided written informed consent before the experiment. Participants were paid for their time, for their performance in the reward learning task as well as memory for the state-to-state sequences (up to £10 based on percent correct performance above chance), and a bonus for performance in the localizer phase target detection task (up to £2).

### Experimental task

We designed our reward learning experiment to investigate the potential role of replay in prospective planning and memory preservation (**Fig. 1a**). We adapted a reward-based learning task used in previous experiments to target model-based decision making (50, 51), itself an adaptation of a common ‘two-step’ task (52). In this version of the task, there is an equivalence between the two alternative top-level start states in a world. This equivalence provides us with the ability to dissociate model-based and model-free behavior. An additional condition is that potential reward points drift across trials at a sufficient rate that allows for model-free and model-based expectations to often differ. We refer to the decision process as ‘planning’. Closely related studies (40, 41, 71) have defined planning as the engagement of model-based decision-making, and our results are consistent with a strong role for model-based decision-making. Of note, and similar to the current design, Miller et al. (41) use the term ‘planning’ in a two-step task where a decision is made solely at the top level.

Additionally, this version utilizes deterministic transitions between states. The deterministic version of the task has been shown to both incentivize and increase model-based behavior (51, 54). A task using probabilistic transitions would have decreased our ability to detect clear evidence of sequential reactivation, given the increase in number of transitions to evaluate and a constrained number of total trials, as reasonable scanning duration was already maximized. This reasoning also led to using a task with a non-branching path structure (i.e. where states involved no forced choices). We also constrained choice points and branching paths in order to decrease the number of total states, which maintained our ability to sufficiently decode visual states using state-of-the-art imaging technology and analysis techniques (53). Importantly, this limitation of branching choices is similar to the environments in the majority of rodent studies of replay, where animals navigate linear tracks with few, if any, subsequent choices (e.g. 2, 27, 30, 33, 35, 36, 39, 44). While these situations do not strictly require sequential evaluation (11), such human and animal designs are able to take advantage of any default tendency to utilize such a neural mechanism. Finally, as measures of model-based behavior are dependent on start state alone, the current task omitted choice at the terminal state, allowing participants to better track rewards and thereby likely increase model-based behavior (41, 51, 54, following recent work; 78).

We augmented this task by adding a second multi-step structure or ‘world’ in order to study the relationship between neural replay signatures and memory preservation (**Fig. 1a**). Further, by employing two worlds we 1) decreased the predictability of the upcoming trial, which in turn increased the utility of planning at choice onset versus immediately after feedback (56), and 2) decreased the dependence of reward learning on short-term working memory for immediately preceding feedback (79, 80). We ensured that participants understood the structure of the task in four ways, as poor understanding of a related paradigm has been reported to result in apparent model-free behavior (69). First, we provided extensive instructions to minimize misunderstanding. Second, we employed an initial non-reinforced structure exploration learning phase. Third, across all sessions, accurate knowledge of task structure itself was incentivized by periodic memory test trials that allowed participants to harvest additional payment. Fourth, participants learned the abstract task structure during an initial training session (with different stimuli). This session was followed by a break before the MEG session, allowing for robust learning of the structure prior to scanning and potentially allowing for memory consolidation. We conducted the first session on a preceding day for almost all participants (n = 22, range 1-13 days) or after a 2-hour break in two participants, resulting in a 2 day median separation between sessions. Overall, these features helped ensure that behavior was quite model-based, as our goal was to understand planning. Further, by studying behavior reliant on well-learned structure knowledge, our results may better connect to how memory is utilized outside the lab.

### Session 1 behavioral procedure

#### Instructions

In the first experimental session, participants were first instructed in the two-world task, then completed a structure learning phase, and finally completed a brief reward learning phase. This session utilized the same abstract structure as the subsequent MEG session but the structures were populated with different visual stimuli.

Participants were given detailed instructions about the reward learning task and the underlying ‘world’ structure in order to maximize understanding. In brief, they were first instructed that on each trial they would face a choice between two different shape options. Each shape would lead to a different path made up of three sequentially-presented images. Paths ended in reward points, which ranged from 0-9, and these reward points would drift over time. Their goal was to choose the shape that led down the path to the currently greater number of reward points to earn more bonus money in the experiment. They were also told that along with the presentation of shapes, they would see either ‘1x’ or ‘5x’ above the shape options, which indicated whether the end reward points on this trial would be multiplied by 1 or by 5 (51).

Next, participants were told that there were actually two pairs of shape options that would converge to lead down the same two paths and reach the same source of rewards. The shapes in the two pairs were equivalent, in that if shape 1a in start state A led to the path with a snowflake, shape 1b in start state B would also lead to the path with the snowflake (**Fig. 1a**). Importantly, participants were instructed that the rewards at the end of the paths would be reached by either of the potential starting shapes (e.g. 1a and 1b). Consequently, if they were first choosing between the shapes in start state A and found a high reward after choosing shape 1a, then if they were subsequently starting with pair B, they should choose shape 1b in order to get to the same source of reward as they had just experienced when they chose shape 1a. Participants can only accomplish this by leveraging their knowledge of the task structure, classifying this as model-based behavior.

Finally, participants were instructed that these shapes and paths made up 1 ‘world’ and that the real experiment would have two independent worlds, each with the same structure. Trials would start with a pair of shapes from one of the two worlds at random. In general, participants’ goal was to keep track of what paths led to the highest rewards at a given time and to choose the shape that led to those paths, while at the same time maintaining their memory for the paths in each world.

#### Structure learning session 1

Before starting the full reward learning task with point feedback, participants were given the opportunity to learn the structure of the worlds. This phase was composed of two blocks of 20 trials, with learning incentivized by rewarding performance on memory questions about the structure of the worlds. Trials started pseudorandomly in one of the four potential start states across the two worlds.

Participants’ goal in this structure learning phase was to explore the different paths in order to learn the sequence of images that followed each shape. Trial events were the same as in reward learning phase trials (see below; **Fig. 1b**; **Fig. S1a**) with the exception that no reward points were shown at the end of the paths and no stakes information was shown during shape presentation. In each structure learning trial, after a planning period, participants made a choice between two shape options (shown randomly on the left and right of the screen). After this selection, the three following states in the path associated with that shape were presented sequentially. Participants were instructed to remember the complete sequence from the chosen shape through to the third path picture. Each path stimulus was randomly presented on the left or right side of the screen and participants needed to press the corresponding 1 or 2 button to indicate the stimulus location on the screen. In this phase, no reward feedback was presented. A fixation cross was presented during a 4-6 s inter-trial interval (ITI). (See detailed timing in the MEG reward learning phase description.)

After each of the two structure learning blocks of 20 trials, participants completed eight memory test probe trials (**Fig. S1b**). Each of the eight start shapes cued one probe trial. On a memory test trial, a single shape was presented on the left or right side of the screen. After the participant selected the shape, they were presented with four potential stimuli from the first state of each of the four paths. Participants selected the stimulus that came next using the 1-4 buttons. After framing the selected stimulus in blue for 0.25 s, this probe structure was repeated for the second and third states in the path. At each stage, one of the four stimuli was correct, while the other three stimuli came from the same state (first, second, or third) across the other three paths. After the response for the third path stimulus, participants were asked to rate their confidence in their set of answers according to the following scale and indicated button response: “Guess (1) Low (2) Medium (3) Certain (4)”. Correct performance was based on accurately selecting the correct picture at each of the three stages. Structure memory performance increased from 56.9% (range, 0-100) after the first 20 learning trials to 90.1% (range, 12.5-100) after the second 20 learning trials. Similarly, mean confidence ratings increased from 2.89 to 3.65.

#### Reward learning session 1

Next, participants engaged in a short reward learning phase to provide experience in maximizing reward earnings. The reward learning phase was the same as the scanned reward learning phase in session 2 (below), with the exception that structure memory questions were pseudo-randomly interleaved with the regular reward learning trials instead of being segregated to breaks between blocks. Trials started pseudorandomly in one of the four potential start states across the two worlds. All trials proceeded in the same way as trials in the preceding structure learning phase, with the addition of reward feedback at the end of each path as well as cued stakes information during planning and choice (**Fig. 1b**; **Fig. S1a**). Reward feedback was presented after a 2 s interstimulus-interval (ISI). Reward points flickered in brightness for a period of 1.5 s and then a 3-5 s blank inter-trial interval followed, a slight difference in procedure from the subsequent MEG session. (See detailed timing in the MEG reward learning phase description.)

The reward learning phase length in session 1 was initially based on the free time remaining in the scheduled session but was then set to be a maximum of 40 trials, resulting in a mean of 50 trials across participants (range, 29-105 trials). The memory probe questions were made more difficult in the reward learning phase than the structure learning phase by randomizing the incorrect lure stimuli to be from any stage and any path. Performance on the interleaved structure memory probes was 92.2% (range 70-100%). As noted above, one participant was not invited for MEG scanning based on very poor session 1 memory probe performance (40% correct).

### Session 2 MEG procedure

#### Localizer

After initial setup in the MEG room, participants were given instructions for the localizer phase. The purpose of the functional localizer was to derive participant-specific sensor patterns that discriminated each of the 12 stimuli that made up the world paths by presenting each stimulus many times. The localizer design was adapted from those used previously, where a picture name identification task followed the presentation of each picture stimulus (11). Participants were instructed to pay attention to a picture shown in the center of the screen and then after the picture disappeared, to select the correct name for the picture from two alternatives. For complete localizer phase details, see **Supplementary Methods**.

#### Structure learning session 2

Participants then engaged in a structure learning phase where the new stimuli from the preceding localizer populated the two worlds. This phase was the same as the no-reward structure learning phase in session 1 (above). Participants were reminded of the world structure. For analyses, accuracy focused on the response for the first transition. We found participants explored all of the eight potential shape-path combinations (most-explored path per-participant, mean 6.8 trials out of 40 total [range 6.0 – 8.4]; least-explored path per-participant, mean 3.3 trials [range 2.0 – 4.2]).

#### Reward learning session 2

Participants then engaged in the primary reward learning phase. The design of this phase was the same as the reward phase in session 1 (**Fig. 1b**; **Fig. S1a**). Participants aimed to earn as many points as possible in the two different worlds. This phase was composed of 6 blocks of 24 trials for MEG data collection, yielding 144 total reward learning trials. See Supplementary Methods for full details.

### Behavioral analysis

Learning behavior was analyzed using computational models, following prior studies (e.g. 50, 51, 52). To verify learning and determine how reward influenced choice, we used computational models that seek to explain a series of choices in terms of previous events. First, we used logistic regression models, which test for local trial-to-trial adjustments in behavior guided by minimal assumptions about their form, yet at the same time qualitatively capture aspects of model-free and model-based behavior (see **Supplementary Methods**). We then used Q-learning reinforcement learning models, which use a structured set of assumptions to capture longer-term coupling between events and choices and can capture model-free and model-based contributions (see **Supplementary Methods**).

### Rarity of experience

To examine potential memory preservation effects after reward feedback, we computed a variable representing the relative inverse frequency (‘rarity’) of the alternative world. Frequency of experience was an exponentially-weighted measure computed for each world on a trial-by-trial basis. This measure was calculated by adapting the fractional updating in Equation 2 to track world frequency based on appearance (1) or non-appearance (0) of a world on a given trial. We expected the learning rate controlling the change in estimates of world frequency to be relatively slow based on related work (81), which led us to set the experience learning rate to 0.10. The resulting experience frequency values were then inverted to provide the infrequency or ‘rarity’ of each world at each trial. A subsequent control analysis made no assumptions about learning rate, simply testing whether other world replay was stronger when that world had been experienced on the last trial or not.

### MEG Pre-processing

For complete details on MEG acquisition and initial pre-processing, see **Supplementary Methods**. For planning analyses, we used the 2.5 s pre-choice planning period after excluding the first 160 ms to allow for early visual stimulus processing, following a related previous experiment (7). For memory analyses in the post-feedback period, where our prediction was that memory processes would be engaged when other cognitive demands are relatively low, with similarities to a procedural step in a related rodents study (26), we focused on the time period following initial reward processing (the latter 3.5 s of the 5 s period). Here we expected that the demands of actual feedback processing and value updating would preclude the engagement of any memory preservation signal. However, we note that our results remain qualitatively the same even without this early time period exclusion step.

### MEG data decoding and cross-validation

Lasso-regularized logistic regression models were trained for each of the 12 stimuli from the paths. Methods followed previous studies (7–9); for additional details see **Supplementary Methods**. Decoding models were trained on MEG data elicited by direct presentations of the visual stimuli. Our experimental task was not optimized to detect reactivation of expected value or reward point outcome representations, and as a consequence our analyses focus only on stimuli.

### Sequenceness measure

The decoding model described above allowed us to measure spontaneous sequential reactivation of the 12 states either during the planning or after feedback periods. We applied each of the 12 trained classifiers to the MEG data at each time point in each period. This yielded a [time x state] reactivation probability matrix for each period in each trial, containing twelve time series of reactivation probabilities each with the length of time samples included in the analysis window. Please see **Supplementary Methods** for complete details.

We then used the Temporally Delayed Linear Modelling framework to quantify evidence of ‘sequenceness’ (53), which describes the degree to which these representations were reactivated in a task-defined sequential order (7–9, 11, 53). TDLM is a multiple linear regression approach that quantifies the degree to which a time-lagged reactivation timecourse of state *j*, (*X*(Δ*t*)_j_, where Δ*t* indicates lag time) predicts the reactivation timecourse of state *i*, (*X_i_*). It involves two stages. At the first stage, we use a set of multiple linear regression models to generate the empirical state-to-state reactivation pattern, using each state’s (*j* ∈ [1: 12]) reactivation timecourse as a dependent variable, and the historical (i.e., time-lagged) reactivation timecourses of all states (*i* ∈ [1: 12]) as predictor variables.

At the second stage, we quantified the evidence that the empirical transition matrix can be predicted by the sequences of interest, i.e., the 4 paths across both worlds in the task. All transitions of interest were specified in model transition matrices, separately for a forward direction (*T_F_*, the same as visual experience) and the inverse for a backward direction (*T_B_*). As control variables, the regression included a constant matrix (*T_cons_*) that captures the average of all transitions, ensuring that any identified effects were not due to background neural dynamics, and a matrix (*T_auto_*) that models self-transitions to control for auto-correlation. Repeating this procedure at each time lag (Δ*t* = 10, 20, 30, &, 600 ms) results in timecourses of both forward and backward sequence strength as a function of time lag, where smaller lags indicate greater time-compression of replay.

Please see **Supplementary Methods** for the following sections: Identifying Replay Onsets, MEG Source Reconstruction, Time-frequency analyses, Non-sequential reactivation analyses, Representational similarity, Multilevel regression models, and Statistical correction and null effects.

## Data availability

Complete behavioral data are publicly available on the Open Science Framework (https://osf.io/szjxp/). The full MEG dataset will be publicly available on openneuro.org.

## Code availability

Example code for sequenceness analyses will be available on GitHub (https://github.com/gewimmer-neuro/multistep_replay).

## Acknowledgments

The authors thank Rani Moran for helpful discussions, Matt Nour for assistance with analysis, and Daniel Bates and the Imaging Support team at WCHN for assistance with scanning. This work was supported by a Wellcome Investigator Award (098362/Z/12/Z) to R.J.D. G.E.W is supported by a research fellowship from the Deutsche Forschungsgemeinschaft (DFG) and an MRC Career Development Award (MR/V032429/1). Y.L. is supported by the Open Research Fund of the State Key Laboratory of Cognitive Neuroscience and Learning. D.M is supported by a Sir Henry Wellcome Trust Postdoctoral Research Fellowship (110257/Z/15/Z). The Max Planck University College London Centre is a joint initiative supported by University College London and the Max Planck Society. The Wellcome Centre for Human Neuroimaging is supported by core funding from the Wellcome Trust (203147/Z/16/Z).

This research was funded in whole, or in part, by the Wellcome Trust 098362/Z/12/Z. For the purpose of Open Access, the author has applied a CC BY public copyright license to any Author Accepted Manuscript version arising from this submission.

## Author contributions

G.E.W., Y.L., and D.C.M. designed the experiment. G.E.W. collected and analyzed the data. G.E.W and Y.L. wrote the analysis code. G.E.W, Y.L., D.C.M and R.D. contributed to data interpretation. G.E.W. wrote the paper with Y.L., D.C.M., and R.J.D.

## References

1. H. F. Olafsdottir, D. Bush, C. Barry, The Role of Hippocampal Replay in Memory and Planning. Current biology: CB 28, R37–R50 (2018).

2. K. Diba, G. Buzsaki, Forward and reverse hippocampal place-cell sequences during ripples. Nat Neurosci 10, 1241–1242 (2007).

3. G. Buzsaki, Hippocampal sharp wave-ripple: A cognitive biomarker for episodic memory and planning. Hippocampus 25, 1073–1188 (2015).

4. H. R. Joo, L. M. Frank, The hippocampal sharp wave-ripple in memory retrieval for immediate use and consolidation. Nat Rev Neurosci 19, 744–757 (2018).

5. D. J. Foster, Replay Comes of Age. Annu Rev Neurosci 40, 581–602 (2017).

6. A. M. Wikenheiser, A. D. Redish, Hippocampal theta sequences reflect current goals. Nat Neurosci 18, 289–294 (2015).

7. G. E. Wimmer, Y. Liu, N. Vehar, T. J. Behrens, R. D. Dolan, Episodic memory retrieval success is related to rapid replay of episode content. Nat Neurosci https://doi.org/10.1038/s41593-020-0649-z (2020).

8. Y. Liu, R. J. Dolan, Z. Kurth-Nelson, T. Behrens, Human replay spontaneously reorganises experience. Cell 178, 640–652 e614 (2019).

9. Z. Kurth-Nelson, M. Economides, R. J. Dolan, P. Dayan, Fast Sequences of Non-spatial State Representations in Humans. Neuron 91, 194–204 (2016).

10. E. Eldar, G. Lievre, P. Dayan, R. J. Dolan, The roles of online and offline replay in planning. Elife 9 (2020).

11. Y. Liu, M. G. Mattar, T. E. J. Behrens, N. D. Daw, R. J. Dolan, Experience replay is associated with efficient nonlocal learning. Science 372 (2021).

12. M. M. Nour, Y. Liu, A. Arumuham, Z. Kurth-Nelson, R. J. Dolan, Impaired neural replay of inferred relationships in schizophrenia. Cell 10.1016/j.cell.2021.06.012 (2021).

13. T. Wise, Y. Z. Liu, F. Chowdhury, R. J. Dolan, Model-based aversive learning in humans is supported by preferential task state reactivation. Science Advances 7 (2021).

14. S. Michelmann, B. P. Staresina, H. Bowman, S. Hanslmayr, Speed of time-compressed forward replay flexibly changes in human episodic memory. Nat Hum Behav 3, 143–154 (2019).

15. G. Pezzulo, F. Donnarumma, D. Maisto, I. Stoianov, Planning at decision time and in the background during spatial navigation. Curr Opin Beh Sci 29, 69–76 (2019).

16. R. J. Dolan, P. Dayan, Goals and habits in the brain. Neuron 80, 312–325 (2013).

17. G. Pezzulo, C. Kemere, M. A. A. van der Meer, Internally generated hippocampal sequences as a vantage point to probe future-oriented cognition. Ann N Y Acad Sci 1396, 144–165 (2017).

18. R. L. Buckner, The role of the hippocampus in prediction and imagination. Annu Rev Psychol 61, 27–48, C21-28 (2010).

19. G. M. van de Ven, H. T. Siegelmann, A. S. Tolias, Brain-inspired replay for continual learning with artificial neural networks. Nat Commun 11, 4069 (2020).

20. J. L. McClelland, B. L. McNaughton, R. C. O’Reilly, Why there are complementary learning systems in the hippocampus and neocortex: insights from the successes and failures of connectionist models of learning and memory. Psychol Rev 102, 419–457 (1995).

21. M. F. Carr, S. P. Jadhav, L. M. Frank, Hippocampal replay in the awake state: a potential substrate for memory consolidation and retrieval. Nat Neurosci 14, 147–153 (2011).

22. D. Kumaran, D. Hassabis, J. L. McClelland, What Learning Systems do Intelligent Agents Need? Complementary Learning Systems Theory Updated. Trends Cogn Sci 20, 512–534 (2016).

23. G. M. van de Ven, S. Trouche, C. G. McNamara, K. Allen, D. Dupret, Hippocampal Offline Reactivation Consolidates Recently Formed Cell Assembly Patterns during Sharp Wave-Ripples. Neuron 92, 968–974 (2016).

24. D. M. McNamee, K. L. Stachenfeld, M. Botvinick, S. J. Gershman, Flexible modulation of sequence generation in the entorhinal-hippocampal system. Nat Neurosci (2021).

25. M. A. Wilson, B. L. McNaughton, Reactivation of hippocampal ensemble memories during sleep. Science 265, 676–679 (1994).

26. H. F. Olafsdottir, F. Carpenter, C. Barry, Task Demands Predict a Dynamic Switch in the Content of Awake Hippocampal Replay. Neuron 96, 925–935 e926 (2017).

27. A. E. Papale, M. C. Zielinski, L. M. Frank, S. P. Jadhav, A. D. Redish, Interplay between Hippocampal Sharp-Wave-Ripple Events and Vicarious Trial and Error Behaviors in Decision Making. Neuron 92, 975–982 (2016).

28. A. S. Gupta, M. A. van der Meer, D. S. Touretzky, A. D. Redish, Hippocampal replay is not a simple function of experience. Neuron 65, 695–705 (2010).

29. A. Johnson, A. D. Redish, Neural ensembles in CA3 transiently encode paths forward of the animal at a decision point. J Neurosci 27, 12176–12189 (2007).

30. R. E. Ambrose, B. E. Pfeiffer, D. J. Foster, Reverse Replay of Hippocampal Place Cells Is Uniquely Modulated by Changing Reward. Neuron 91, 1124–1136 (2016).

31. B. E. Pfeiffer, D. J. Foster, Hippocampal place-cell sequences depict future paths to remembered goals. Nature 497, 74–79 (2013).

32. H. F. Olafsdottir, C. Barry, A. B. Saleem, D. Hassabis, H. J. Spiers, Hippocampal place cells construct reward related sequences through unexplored space. Elife 4, e06063 (2015).

33. J. D. Shin, W. Tang, S. P. Jadhav, Dynamics of Awake Hippocampal-Prefrontal Replay for Spatial Learning and Memory-Guided Decision Making. Neuron 104, 1110–1125 e1117 (2019).

34. M. G. Mattar, N. D. Daw, Prioritized memory access explains planning and hippocampal replay. Nat Neurosci 21, 1609–1617 (2018).

35. H. Xu, P. Baracskay, J. O’Neill, J. Csicsvari, Assembly Responses of Hippocampal CA1 Place Cells Predict Learned Behavior in Goal-Directed Spatial Tasks on the Radial Eight-Arm Maze. Neuron 101, 119–132 e114 (2019).

36. C. T. Wu, D. Haggerty, C. Kemere, D. Ji, Hippocampal awake replay in fear memory retrieval. Nat Neurosci 20, 571–580 (2017).

37. A. C. Singer, M. F. Carr, M. P. Karlsson, L. M. Frank, Hippocampal SWR activity predicts correct decisions during the initial learning of an alternation task. Neuron 77, 1163–1173 (2013).

38. A. K. Gillespie et al., Hippocampal replay reflects specific past experiences rather than a plan for subsequent choice. Neuron 10.1016/j.neuron.2021.07.029 (2021).

39. A. A. Carey, Y. Tanaka, M. A. A. van der Meer, Reward revaluation biases hippocampal replay content away from the preferred outcome. Nat Neurosci 22, 1450–1459 (2019).

40. O. M. Vikbladh et al., Hippocampal contributions to model-based planning and spatial memory. Neuron 102, 683–693 e684 (2019).

41. K. J. Miller, M. M. Botvinick, C. D. Brody, Dorsal hippocampus contributes to model-based planning. Nat Neurosci 20, 1269–1276 (2017).

42. R. Ratcliff, Connectionist models of recognition memory: constraints imposed by learning and forgetting functions. Psychol Rev 97, 285–308 (1990).

43. M. McCloskey, N. J. Cohen, Catastrophic interference in connectionist networks: the sequential learning problem. Psychol Learn Motiv 24, 109–165 (1989).

44. S. P. Jadhav, C. Kemere, P. W. German, L. M. Frank, Awake hippocampal sharp-wave ripples support spatial memory. Science 336, 1454–1458 (2012).

45. G. Girardeau, K. Benchenane, S. I. Wiener, G. Buzsaki, M. B. Zugaro, Selective suppression of hippocampal ripples impairs spatial memory. Nat Neurosci 12, 1222–1223 (2009).

46. V. Ego-Stengel, M. A. Wilson, Disruption of ripple-associated hippocampal activity during rest impairs spatial learning in the rat. Hippocampus 20, 1–10 (2010).

47. A. Fernandez-Ruiz et al., Long-duration hippocampal sharp wave ripples improve memory. Science 364, 1082–1086 (2019).

48. N. W. Schuck, Y. Niv, Sequential replay of nonspatial task states in the human hippocampus. Science 364 (2019).

49. A. C. Schapiro, E. A. McDevitt, T. T. Rogers, S. C. Mednick, K. A. Norman, Human hippocampal replay during rest prioritizes weakly learned information and predicts memory performance. Nat Commun 9, 3920 (2018).

50. B. B. Doll, K. D. Duncan, D. A. Simon, D. Shohamy, N. D. Daw, Model-based choices involve prospective neural activity. Nat Neurosci 18, 767–772 (2015).

51. W. Kool, F. A. Cushman, S. J. Gershman, When Does Model-Based Control Pay Off? PLoS Comput Biol 12, e1005090 (2016).

52. N. D. Daw, S. J. Gershman, B. Seymour, P. Dayan, R. J. Dolan, Model-based influences on humans’ choices and striatal prediction errors. Neuron 69, 1204–1215 (2011).

53. Y. Liu et al., Temporally delayed linear modelling (TDLM) measures replay in both animals and humans. Elife 10 (2021).

54. W. Kool, S. J. Gershman, F. A. Cushman, Cost-Benefit Arbitration Between Multiple Reinforcement-Learning Systems. Psychol Sci 28, 1321–1333 (2017).

55. E. H. Patzelt, W. Kool, A. J. Millner, S. J. Gershman, Incentives Boost Model-Based Control Across a Range of Severity on Several Psychiatric Constructs. Biol Psychiatry 85, 425–433 (2019).

56. A. Konovalov, I. Krajbich, Gaze data reveal distinct choice processes underlying model-based and model-free reinforcement learning. Nat Commun 7, 12438 (2016).

57. D. Lakens, Equivalence Tests: A Practical Primer for t Tests, Correlations, and Meta-Analyses. Soc Psychol Personal Sci 8, 355–362 (2017).

58. T. J. Davidson, F. Kloosterman, M. A. Wilson, Hippocampal replay of extended experience. Neuron 63, 497–507 (2009).

59. G. E. Wimmer, N. D. Daw, D. Shohamy, Generalization of value in reinforcement learning by humans. Eur J Neurosci 35, 1092–1104 (2012).

60. M. Lebreton, S. Jorge, V. Michel, B. Thirion, M. Pessiglione, An automatic valuation system in the human brain: evidence from functional neuroimaging. Neuron 64, 431–439 (2009).

61. D. Kumaran, J. J. Summerfield, D. Hassabis, E. A. Maguire, Tracking the emergence of conceptual knowledge during human decision making. Neuron 63, 889–901 (2009).

62. A. Lopez-Persem et al., Four core properties of the human brain valuation system demonstrated in intracranial signals. Nat Neurosci 23, 664–675 (2020).

63. C. Higgins et al., Replay bursts in humans coincide with activation of the default mode and parietal alpha networks. Neuron 109, 882–893 e887 (2021).

64. C. Sun, W. Yang, J. Martin, S. Tonegawa, Hippocampal neurons represent events as transferable units of experience. Nat Neurosci 23, 651–663 (2020).

65. J. Diedrichsen, N. Kriegeskorte, Representational models: A common framework for understanding encoding, pattern-component, and representational-similarity analysis. PLoS Comput Biol 13, e1005508 (2017).

66. L. Deuker, J. L. Bellmund, T. Navarro Schroder, C. F. Doeller, An event map of memory space in the hippocampus. Elife 5 (2016).

67. M. Sosa, H. R. Joo, L. M. Frank, Dorsal and Ventral Hippocampal Sharp-Wave Ripples Activate Distinct Nucleus Accumbens Networks. Neuron 105, 725–741 e728 (2020).

68. S. N. Gomperts, F. Kloosterman, M. A. Wilson, VTA neurons coordinate with the hippocampal reactivation of spatial experience. Elife 4 (2015).

69. C. Feher da Silva, T. A. Hare, Humans primarily use model-based inference in the two-stage task. Nat Hum Behav 10.1038/s41562-020-0905-y (2020).

70. S. Kali, P. Dayan, Off-line replay maintains declarative memories in a model of hippocampal-neocortical interactions. Nat Neurosci 7, 286–294 (2004).

71. A. R. Otto, S. J. Gershman, A. B. Markman, N. D. Daw, The curse of planning: dissecting multiple reinforcement-learning systems by taxing the central executive. Psychol Sci 24, 751–761 (2013).

72. T. E. J. Behrens et al., What Is a Cognitive Map? Organizing Knowledge for Flexible Behavior. Neuron 100, 490–509 (2018).

73. C. R. Brewin, J. D. Gregory, M. Lipton, N. Burgess, Intrusive images in psychological disorders: characteristics, neural mechanisms, and treatment implications. Psychol Rev 117, 210–232 (2010).

74. J. Suh, D. J. Foster, H. Davoudi, M. A. Wilson, S. Tonegawa, Impaired hippocampal ripple-associated replay in a mouse model of schizophrenia. Neuron 80, 484–493 (2013).

75. Q. J. M. Huys, M. Browning, M. P. Paulus, M. J. Frank, Advances in the computational understanding of mental illness. Neuropsychopharmacology 46, 3–19 (2021).

76. E. A. Jones, A. K. Gillespie, S. Y. Yoon, L. M. Frank, Y. Huang, Early Hippocampal Sharp-Wave Ripple Deficits Predict Later Learning and Memory Impairments in an Alzheimer’s Disease Mouse Model. Cell Rep 29, 2123–2133 e2124 (2019).

77. S. M. Prince et al., Alzheimer’s pathology causes impaired inhibitory connections and reactivation of spatial codes during spatial navigation. Cell Rep 35, 109008 (2021).

78. C. M. Gillan, A. R. Otto, E. A. Phelps, N. D. Daw, Model-based learning protects against forming habits. Cogn Affect Behav Neurosci 10.3758/s13415-015-0347-6 (2015).

79. A. G. Collins, M. J. Frank, How much of reinforcement learning is working memory, not reinforcement learning? A behavioral, computational, and neurogenetic analysis. Eur J Neurosci 35, 1024–1035 (2012).

80. G. E. Wimmer, J. K. Li, K. J. Gorgolewski, R. A. Poldrack, Reward learning over weeks versus minutes increases the neural representation of value in the human brain. J Neurosci 38, 7649–7666 (2018).

81. A. M. Bornstein, N. D. Daw, Cortical and hippocampal correlates of deliberation during model-based decisions for rewards in humans. PLoS Comput Biol 9, e1003387 (2013).

