## Supplementary Information for "Distinct replay signatures for prospective decision-making and memory preservation"

**This PDF file includes:**

Supplementary Methods
Supplementary Text
Tables S1 to S4
Figures S1 to S9
SI References

### Supporting methods

#### Behavioral procedure

#### Localizer phase details

The instructions were followed by 12 practice trials. At the mid-point of each localizer block, a 20 s rest was inserted (after which participants pressed a key to continue). After each block, performance was indicated by 1-4 yellow stars indicating performance >65% correct through >95% correct. If performance was lower than 65%, participants were shown the text: "Please try to perform better in the next part!"

In a localizer trial, participants were presented with a stimulus in the center of the screen for 0.85 s. Then the stimulus disappeared and two words appeared, one above the screen midpoint and the other below. One of the words correctly named the preceding object, while the other was incorrect; the top / bottom locations of the correct and incorrect word were randomized. Participants were instructed to select the correct word using the $1^{st}$ or $2^{nd}$ buttons on a 4-button response pad, which was rotated in orientation such that the $1^{st}$ button was above the $2^{nd}$. Words were presented for 0.45 s, followed by a fixation ITI during which button responses for the words were still recorded. The ITI duration mean was 1.2 s (range 0.7 – 2.7 s). If performance on the naming task fell below 70% correct (across missed responses and false alarms in the preceding 32 trials), a warning was presented: "Please increase your performance on the picture identification!"

The stimulus images were presented in a pseudo-random order, with the constraint that no stimulus repeat in subsequent trials. Each stimulus was presented 50 times. The localizer was divided into 4 blocks, with 150 trials per block for a total of 600 trials.

#### Reward learning session 2 details

A reward learning trial began with a 2.5 s planning period where a response was not allowed. To indicate this period, a black cross was presented in the center of the screen. Above the two shape options, the reward stakes on the current trial (1x or 5x) were shown. When the fixation cross disappeared, participants could select a shape for a maximum of 1.5 s. Note that this enforced delay before choice execution limits the ability to interrogate choice reaction time. If no choice response was recorded for this choice, the trial ended with text indicating a loss of nine points: “Too late or wrong key! -9 pts”. After a 0.5 s inter-stimulus interval (ISI) the stimulus from the first path state was presented randomly on the left or right side of the screen for a minimum of 0.5 s up until a response was recorded, or 1.5 s max. Participants responded with a button corresponding to the screen location of the stimulus. After a 0.5 s ISI, this procedure was repeated for the second and third path state stimuli. If participants failed to respond to any path stimuli, the trial ended with the above no-response error message. Either a 0.5 s or 7.0 s ISI preceded the feedback period. This pre-feedback delay in one world (randomly assigned) was always 0.5 s while the delay in the other world was 7.0 s. (This delay had no effect on behavioral or neural results and thus all analyses combine results across worlds.) The feedback points were then presented in colored text determined by the point amount, where a bright green color was used for the maximum amount of 9 points and bright red color for the minimum amount of 0 points (with intermediate point values colored along a green-to-red gradient). Below the display of the points, the total point value accounting for the stakes multiplier was shown. The feedback text initially flickered in brightness for 0.75 s. Next, the text faded away across a period of 2.25 s after which the screen was blank for the remaining ITI of 2-3 s, for a total feedback period of 5-6 s.

We generated a set of counterbalanced lists in order to decrease variability between participants. Trial lists and counterbalancing assignments for the task were generated for reward point drifts, world order, start state order within world, stimulus assignments to states, shape assignment to paths, and stakes. First, reward points drifted from trial-to-trial according to a process using a standard deviation of 1.75 and reflecting boundaries of 0 and 9. Additional criteria included a mean point value between 4.45 and 4.55 for each path in each half of the task and a weak negative correlation between reward points in a given world of between -0.175 and -0.225. Two pairs of reward point drifts of 72 trials were generated, corresponding to the two worlds. One of the paths in each world was initialized with a start value of 8 while the other was initialized with a start value of 2. The reward point drifts were counterbalanced across worlds across participants. Second, world order was pseudorandomized with predominant alternation across trials and a maximum repetition of the same world of 3 trials. Third, within a given world, the start state was pseudorandomized with predominant alternation of start state across trials and a maximum repetition of the same start state in a world of 4 trials. Fourth, stimuli were assigned to states based on four counterbalance lists. Fifth, shapes were assigned to paths based on five counterbalance lists. Finally, per-trial stakes were pseudorandomly assigned with a maximum repetition of the same stakes value of 4 trials. For the cued reward stakes manipulation, unfortunately, we identified an unintended significant correlation between stakes and estimated option values (**Supp. Results**). Because of this limitation, we could not clearly assess any relationships between stakes and behavior or neural activity. Further, as we found no benefit from including stakes information in the learning models (**Supp. Results**), the effect of stakes was not included in any behavioral or MEG analyses; i.e. any analyses based on reward points used the unmodulated point value.

At the end of each block of 24 learning trials, participants engaged in a set of memory probe trials during a break in MEG data acquisition. The first two learning blocks were each followed by 8 memory probe questions to ensure the existence of robust structure knowledge, while the remaining four blocks were followed by 5 memory probe questions. As in session 1, the memory probe questions were made more difficult in the reward learning phase than the structure learning phase by randomizing the incorrect lure stimuli to be from any stage and any path. (In the first two participants, confidence ratings were not collected.) See the preceding description in session 1 for memory probe trial timing information. After the memory questions, participants could rest and stretch until ready for the next block.

In the first two participants, only five learning blocks were collected and the learning list also varied, resulting in different numbers of trials with MEG data (100 and 88). In these participants, the memory test questions were randomly interspersed between choice trials instead of being segregated into mini-blocks. In the third participant, only five learning blocks were collected, leading to 120 trials with MEG data. All analyses were adjusted to account for the difference in total trials.

For 21 participants, an additional localizer scan was collected after the reward learning blocks; these data are not analyzed here. Following scanning, all participants then completed a brief written post-experiment questionnaire.

**Behavioral analysis**

Primary analyses examined behavior independent of whether a trial (or trials) from the alternative world were interleaved between current world trials. All missed response trials (where no response was recorded within the response time window; mean = 3.75 trials per participant) were excluded from the below analyses. In the stay and switch analyses, the previous non-missed trial was counted as the last choice. For reinforcement learning models, the Q-value estimates were carried forward, ignoring missed trials.

**Regression**

In the regression analysis, we used logistic regression to account for each participant's sequence of choices. For all regression analyses, we focus on the influence of preceding events within the same world (ignoring intervening trials related to the other world). We used multilevel regression functions implemented in R (lme from the nlme package for linear regression; glmmTMB from the glmmTMB package for logistic regression). All predictors and interactions were included as random effects, following the 'maximal' approach (Barr et al., 2013). Correlations between random effects were included when convergence was achievable with this structure.

The first model predicted stay choices based on previous reward and whether the current start state was the same or different (Kool et al., 2016). Here, if participants receive a relatively high reward on the preceding trial, it is generally advantageous in the current task to stay with that choice on the current trial. Conversely, if participants receive a relatively low reward, it is generally advantageous to switch to choosing the alternative option. Thus, the strength of this reward effect on stay choices indexes how well participants were guided by preceding reward. Critically, to test whether experience in one start state is carried over to influence actions in the other start state, our model included a binary variable representing same versus different start state. Any generalization was captured in our model by including the interaction between the previous reward variable and the same start state variable. This measures to what degree, if any, an individual is more influenced by preceding reward when a trial begins in the same start state as the previous trial. Such an influence is characteristic of model-free behavior. Alternatively, given that generalizing across equivalent start states requires knowledge of the structure of the task, the lack of an interaction supports the existence of model-based behavior.

A second model examined how option selection was influenced by the choice and reward received on the previous trial. Instead of predicting stay decisions, this model predicts option choice (with options arbitrarily coded as 0 and 1) using the history of preceding rewards and option choices (Lau and Glimcher, 2005; Wimmer et al., 2014; Doll et al., 2015). (Such models approximate reinforcement learning models when longer histories of previous choices and rewards are included, with the decaying influence of previous events reflecting learning rate.) Critically, in addition to preceding choices and rewards, to assess model-based behavior, the model also included a variable representing whether the current start state was the same or different than the preceding trial. As in the first analysis, the interaction of this same start state variable with previous trial reward and choice is able to capture any model-free influence on behavior (Doll et al., 2015). As above, if there is a stronger influence of previous events on choice when the start state is the same, this provides evidence for a model-free component to behavior. Conversely, the lack of such an interaction supports model-based behavior.

Supplemental regression analyses were conducted within-participants to analyze individual differences in model-based generalization; here, individual participant fits were derived from the bayesglm function in the arm package to constrain extreme coefficients in one participant.

**Reinforcement learning**

In the reinforcement learning analysis, we fit three different Q-learning reinforcement learning (RL) models to subjects choice behavior (Sutton and Barto, 1998): model-free, model-based, and hybrid models. These models generated participant-specific model parameters and model fit values (log likelihood). These models also allow us to compute trial-by-trial variables related to choice value and reward feedback for use in neural analyses.

The model-free reinforcement learning model learns to assign an independent action value to each state of the task. For simplicity, we assign model-free values to the path as a whole, without considering the value of each intermediate state (see discussion below), yielding two "states" in the experiment: the choice state (state 1) and the path (state 2). In this way, the model below is equivalent to and based on the model applied to two-step tasks lacking the intermediate path states (Kool et al., 2016). Further, with no second stage choice, there are only two actions, $a_1$ and $a_{2,}$, with one action selected in the start state. Model-free values are updated at stage *i* and trial *t* according to a prediction error δ, modulated by a learning rate α (with range [0 1]):

$$Q_{MF}(s,a) = Q_{MF}(s,a) + \alpha\,\delta_{i,t} \quad (1)$$

For the model-free strategy, the prediction errors differ after moving from the start states (A and B) to one of the paths (states X1 or X2), and from a path to a reward. The values of the start state options are first updated according to the difference in value between the start and path states:

$$\delta_{1,t} = Q_{MF}(s_{X,t},a_{1,t}) - Q_{MF}(s_{1,t},a_{1,t}) \quad (2)$$

After reward feedback (re-scaled for modeling to a range of [0 1]), the values for the path states are updated according to the difference in value between the path state and the reward received:

$$\delta_{2,t} = r_{2,t} - Q_{MF}(s_{X,t}, a_{1,t}) \quad (3)$$

Additionally, the model-free values for the start states are updated after the received reward, modulated by a fractional eligibility trace parameter $e$ (with range [0 1]):

$$\delta_{1,t} = e(\ r_{X,t} - Q_{MF}(s_{X,t},a_{1,t})\ ) \quad (4)$$

This allows for start states to be updated by the reward received after the second state.

The model-based strategy utilizes the learned path (state 2) model-free Q-values in combination with the learned transition matrix between task states.

$$Q_{MB}(s_A,a_j) = P(s_X \mid s_A,a_j)\ Q_{MF}(s_{X1}) + P(s_X \mid s_B,a_j)\ Q_{MF}(s_{X2}) \quad (5)$$

The hybrid reinforcement learning model combines model-free and model-based learning models and computes per-trial option values based on a weighted sum of the two estimates. Our model was based on that used by Kool et al. (2016), where the values input to the softmax choice rule are calculated according to:

$$Q_{net}(s_A,a_j) = \omega Q_{MB}(s_A,a_j) + (1 - \omega)Q_{MF}(s_A,a_j) \quad (6)$$

Given value estimates on a particular trial where participants were choosing between two options, participants are assumed to stochastically with probabilities according to a softmax distribution (Daw et al., 2006):

$$P(s_{i,t}) = \exp(\beta(Q_{net}\ s_{i,t},a) / \textstyle\sum_{a'}(\ \exp(\beta(Q_{net}\ s_{i,t},a')\ ) \quad (7)$$

The free parameter β represents the softmax inverse temperature (with range [0 Inf]), which controls the exclusivity with which choices are focused on the highest-valued option. This exclusivity can reflect certainty in option estimates. Note that since the softmax is also the link function for the logistic regression model discussed above, this analysis also has the form of a regression from Q values onto choices except here, rather than as linear effects, the past rewards enter via the recursive learning of Q, controlled, in nonlinear fashion, by the learning rate parameters.

The purely model-based RL model is a reduced form of the hybrid model where the weighting parameter $\omega$ = 1 and the eligibility parameter $e$ have no effect and are dropped. Conversely, the purely model-free RL model is a reduced form of the hybrid model where $\omega$ = 0.

We found that models with decaying ("forgetting") Q-values for non-experienced states provided a better fit. Specifically, one parameter decayed the value of non-chosen Q values. In the task, some decay of non-chosen Q values is rational, as the reward values drift for both chosen and non-chosen terminal states. A second parameter decayed Q-values for the non-experienced world. The decay parameters were constrained to the range [0 1]. The parameters decayed values to the median reward value of 0.5. Thus, if the non-experienced world decay parameter was less than 1, when both Q-values were above or below 0.5, both values moved toward the median while the distance between values was maintained. In the situation where Q-values were above and below the median, the distance between values decreased.

Thus, on each trial, the value for the non-chosen Q-values was decayed according to the fractional parameter $\tau_{ALT}$:

$$Q_{MF}(s_i,a_{nonchosen}) = (\ (Q_{MF}(s_i,a_{nonchosen}) - 0.5) * \tau_{ALT}) + 0.5 \quad (8)$$

Separately, on each trial, all values in the non-experienced world were decayed according to the fractional parameter $\tau_W$:

$$Q_{MF}(s_i,a_j) = (\ Q_{MF}\ (s_i,a_j) - 0.5) * \tau_W) + 0.5 \quad (9)$$

Parameters were optimized for each subject using an optimization routine based on the fmincon in Matlab. Option values were initialized with mid-range values of 0.5, which also provided the best fit to the data.

To generate per-participant, per-trial variables based on the reinforcement learning models, the models were simulated on each participant's sequence of experiences with their best-fit parameters.

**Reinforcement learning with feedback replay**

To test if feedback period other world replay was directly related to the quality of subsequent choice behavior, providing additional evidence of a role in memory preservation, we estimated an additional reinforcement learning model augmented by neural replay. Enhanced memory preservation was expected to benefit the representations of the other world structure and associated values, which could be related to decreased uncertainty on subsequent choices. Conversely, memory decay was expected to be associated with increased uncertainty. To capture a preservation-related decrease in uncertainty in our reinforcement learning model, we focused on the softmax inverse temperature associated with the translation of learned values into choice. While this parameter is often interpreted as a signal reflecting noise or uncertainty specific to choice processes, this parameter can also reflect longer-term shifts in the uncertainty of underlying memory representations. In our best fit model-based RL model, the uncertainty at choice is over the prospective value representations, where the effect of uncertainty is a function of the uncertainty of memory for 1) the links between the shape options and the paths, 2) the paths and terminal values, and 3) the terminal values themselves. The softmax inverse temperature can be viewed as effectively controlling the width of the distribution of the estimates of the terminal values. With a high inverse temperature, uncertainty (the width of the distribution) is low and choices are strongly influenced by the stored value estimates. Conversely, with a low inverse temperature, choices become more random and undirected. We predicted that transient increases in memory preservation would be associated with decreased uncertainty, captured by a higher inverse temperature.

We extracted trial-by-trial sequenceness values from the preceding feedback period and the planning period. The modulation of uncertainty applied to trials where the 'world' changed from a preceding trial (e.g. from world 1 to world 2, or vice versa). It is only on 'world change' trials where preceding "other world" backward replay represents the path options that are under consideration in the current trial. To determine trials with relatively high backward replay, the sequenceness data were thresholded at the 60th percentile to ensure sufficient choice trials for RL parameter estimation. The threshold was determined by stepping down until the inclusion of at least 10 trials per participant where there was strong replay (minimum number of trials, 11; median, 27). For the replay strength variable, a 1 represented the presence of strong replay and 0 represented the absence. In control analyses, we also examined alternative replay variables derived from the preceding feedback phase as well as the current planning period.

The RL memory preservation model was built upon the above RL model, augmented with two additional softmax inverse temperature parameters. The baseline β applied to trials where the world was the same as the previous trial. $\beta_{wchange}$ applied to trials where the world changed from the previous trial but where there was no strong replay. $\beta_{wchange\_replay}$ applied to trials where the world changed from the previous trial and there was strong replay. We predicted that $\beta_{wchange\_replay}$ would be higher than $\beta_{wchange}$. The choice equations for the augmented model are as follows. For world-same trials, choice was determined by Eq. 7:

World-change trials where $replay_{t-1} = 0$ (low):

$$P(s_{i,t}) = \exp(\beta_{wchange}(Q_{net}\ s_{i,t},a) \,/\, \textstyle\sum_{a'} (\ \exp(\beta_{wchange}(Q_{net}\ s_{i,t},a')\ ) \qquad (10)$$

World-change trials where $replay_{t-1} = 1$ (high):

$$P(s_{i,t}) = \exp(\beta_{wchange\_replay}(Q_{net}\ s_{i,t},a) \,/\, \textstyle\sum_{a'} (\ \exp(\beta_{wchange\_replay}(Q_{net}\ s_{i,t},a')\ ) \qquad (11)$$

**MEG methods**

**MEG acquisition**

The participants were scanned while sitting upright inside an MEG scanner located at the Wellcome Centre for Human Neuroimaging at UCL. A whole-head axial gradiometer

MEG system operating at third order synthetic gradiometry configuration (CTF Omega, VSM MedTech) recorded data continuously at 1200 samples per second, utilizing 272 channels (3 channels of the original 275 channels are not included due to excessive noise in routine testing). Three head position indicator coils were used to locate the position of the participant's head in the three-dimensional space with respect to the MEG sensor array. They were placed on the three fiducial points: the nasion and left and right pre-auricular areas. The coils generate a small magnetic field which is used to localize the head and enable continuous movement tracking. We also used an Eyelink eye-tracking system to record participant's eye movements and blinks; the MEG implementation uses no additional head restraint or support. Eye-tracking was calibrated prior to scanning and data was stored alongside the MEG data in three additional channels at the MEG sampling rate. The task was projected onto a screen suspended in front of the participants. The participants responded during the task using a 4-button response pad to provide their answers (Current Designs), responding with self-selected digits to the first and second buttons. Trial events were written into the MEG data via a TTL parallel port. Event timing was verified by concurrent photodiode recording of trial-event-associated brightness changes in a bottom-left location of the projected screen (not visible to participants); this channel was also written into the MEG data.

**MEG Pre-processing**

MEG data were processed in MATLAB using the packages SPM12 (Wellcome Trust Centre for Neuroimaging) and FieldTrip, following previous procedures (Liu et al., 2019; Wimmer et al., 2020). See https://github.com/gewimmer-neuro/multistep_replay for an example preprocessing script. The CTF data were imported using the OSL package (the OHBA Software Library, from OHBA Analysis Group, OHBA, Oxford, UK). Slow drift was removed by applying a first-order IIR high-pass filter at 0.5 Hz. The data were down-sampled (including anti-aliasing low-pass filter) from 1200 Hz to 100 Hz (for sequenceness analyses, yielding 10 ms per sample) or 400 Hz (for time-frequency analyses) for improved signal to noise ratio and to conserve processing time.

Preprocessing was conducted separately for each block. An initial preprocessing step in OSL identified potential bad channels whose signal characteristics fell outside the normal distribution of values for all sensors. Then independent component analysis (FastICA, http://research.ics.aalto.fi/ica/fastica) was used to decompose the sensor data for each session into 150 temporally independent components and associated sensor topographies. Data from the three eye-tracking channels was also included and flagged for artifact monitoring. Artifact components were classified by automated inspection of the combined spatial topography, timecourse, kurtosis of the timecourse, and frequency spectrum for all components. For example, eye-blink artifacts exhibit high kurtosis (>20), a repeated pattern in the timecourse and consistent spatial topographies. Mains interference has extremely low kurtosis and a frequency spectrum dominated by 50 Hz line noise. Artifacts were then removed by subtracting them out of the data. All subsequent analyses were performed directly on the filtered, cleaned MEG signal from each sensor, in units of femtotesla.

The MEG data were epoched to extract data for each trial. To allow participants to shift from a goal of structure learning to a goal of value-directed choice, the first 4 trials of the reward learning task were excluded from sequenceness analyses; this exclusion had no qualitative effect on the results. In the localizer blocks, a 2.5 s epoch was extracted in each trial, encompassing 0.5 s preceding stimulus onset and continuing past the stimulus and word response. In the pre-choice planning period, we extracted epochs of 2.5 s at the beginning of each trial. In the post-feedback reward period we extracted epochs of 3.5 s, following the first 1.5 s after reward. At this stage, in the epoched data, we further excluded time periods within individual channels that exhibited extreme outlier events in a given trial epoch (defined by values > 7x the mean absolute deviation) by replacing these values with zero.

In the initial scanned structure learning phase, only 40 trials were presented. To increase power, our sequenceness analyses included all valid paths in the current and other worlds. The contrast of late versus early sequenceness subtracted the mean across the first 20 trials from the second 20 trials. In this phase, the participants' goal was exploration and learning; no reward feedback was presented. Thus, the inter-trial-interval was analogous to the feedback period in the reward learning phase. For MEG analyses, we examined the 4 s inter-trial-interval period beginning at the offset of the third and final path stimulus.

**MEG data decoding and cross-validation**

For each stimulus we trained one binomial classifier. Positive examples for the classifier were trials on which that stimulus was presented. Negative examples consisted of two kinds of data: trials when another stimulus was presented, and data from the fixation period preceding stimulus onset. The null data were included to reduce the correlation between different classifiers – enabling all classifiers to report low probabilities simultaneously. Only the sensors that were not rejected across all scanning sessions in preprocessing were used to train the decoding models for a given participant. A trained model k consisted of a single vector with length of good sensors n consisting of 1 slope coefficient for each of the sensors together with an intercept coefficient.

Prediction accuracy was estimated by treating the highest probability output among all classifiers as the predicted stimulus. In the functional localizer task, prediction performance of classifiers trained at each 10 ms bin from 0 ms to 800 ms after stimulus onset and performance was tested iteratively on left-out trials. The range of peak classifier performance across participants was then identified, with the 200 ms time period selected among the peak times for all subsequent analyses based on previous replay experiments (see Results) (Kurth-Nelson et al., 2016; Liu et al., 2019; Wimmer et al., 2020). A new classifier trained at this time point on all the localizer data trials was estimated for use in subsequent decoding analyses. To ensure that the classification results were not overfit to the regularization parameter of the logistic regression, all results were obtained with the lasso regularization parameter that yielded the strongest mean cross-decoding from the localizer to the sequential presentation of the actual stimuli in the reward learning phase. This independent cross-decoding approach yielded the same parameter value (l1 = 0.002) as we found previously (Wimmer et al., 2020).

In a control analysis, prior to the sequential analyses, we examined whether we could find characteristics suggestive of single stimulus reactivation in the reward learning task periods of interest. This control is supplemental, as we do not identify specific events in any of our analyses and the features of the human MEG signal are best analyzed by regression analyses using all the data in periods of interest (Liu et al., 2021a). For this reactivation analysis, we compared the 12 stimulus time series derived from applying the trained stimulus classifiers to the trial-wise planning and feedback period data (when no stimuli were actually being presented) to time series derived from applying permuted classifiers. To create permuted classifier data, within each participant and for each of the 12 trained stimulus classifiers, we took the original trained sensor weights (a 1-dimensional vector) and randomized the vector weightings across the non-zero-weighted sensors, which preserves the use of informative sensors.

We compared two measures of the resulting time series: mean and skewness. In simulations, when true spike-like events were added to a random vector, the positive spike deviations increased the mean signal, and – independent of the mean – the spike events also increased skewness. Comparing our real data to the permuted data, we indeed found significantly higher mean values in the real versus permuted data (averaged across all stimuli, as we found equal reactivation for current and other word stimuli; p-values < 0.01). Second, we found significantly increased skewness in the real versus permuted data (p-values < 0.01). These effects were found in both the planning period and the feedback period; note that in these periods none of the relevant trained stimuli were presented on-screen.

**Sequenceness measure**

Our analysis of sequential reactivation uses Temporally Delayed Linear Modelling (TDLM), as fully described in Liu et al. (Liu et al., 2021a). The decoding model described above allowed us to measure spontaneous sequential reactivation of the 12 states either during the planning or after feedback periods. We applied each of the 12 trained classifiers to the MEG data at each time point in each period. This yielded a [time x state] reactivation probability matrix for each period in each trial, containing twelve time series of reactivation probabilities each with the length of time samples included in the analysis window (**Fig. S3a**).

We then used TDLM to quantify evidence of 'sequenceness', which describes the degree to which these representations were reactivated in a task-defined sequential order (Kurth-Nelson et al., 2016; Liu et al., 2019; Wimmer et al., 2020; Liu et al., 2021a;

Liu et al., 2021b). TDLM is a multiple linear regression approach that quantifies the degree to which a time-lagged reactivation timecourse of state $j$, ($X(\Delta t)_j$, where $\Delta t$ indicates lag time) predicts the reactivation timecourse of state $i$, ($X_i$). It involves two stages. At the first stage, we use a set of multiple linear regression models to generate the empirical state-to-state reactivation pattern, for each period in each trial, using each state's ($i \in [1{:}\ 12]$) reactivation timecourse as a dependent variable, and the historical (i.e., time-lagged) reactivation timecourses of all states ($j \in [1{:}\ 12]$) as predictor variables. Separate linear models were estimated for each stimulus *i* and each time lag $\Delta t$:

$$X_i = \sum_{j=1}^{12} X(\Delta t)_j \beta(\Delta t)_{ji} + C \qquad (12)$$

where *C* is a constant term and $\beta(\Delta t)_{ji}$ is the coefficient capturing the unique variance in $X_i$ explained by $X(\Delta t)_j$. The resulting regression coefficients quantify the evidence for each empirical state -> state reactivation pattern at a specific lag, $\Delta t$. We calculated sequenceness from time lag 10 ms to 600 ms. All such first-level coefficients are represented in a lag-specific empirical transition matrix $\boldsymbol{B}$, representing evidence for all possible state-to-state transitions at a given time lag (**Fig. S3b**).

At the second stage, we quantified the evidence that the empirical transition matrix $\boldsymbol{B}$ can be predicted by the sequences of interest, i.e., the 4 paths across both worlds in the task. All transitions of interest were specified in model transition matrices (of size 12x12, with 1s for transitions of interest and 0s otherwise), separately for a forward direction ($T_F$, the same as visual experience) and the inverse for a backward direction ($T_B$), where $T_F = T_B'$. As control variables, the model included a constant matrix ($T_{cons}$) that captures the average of all transitions (estimating the intercept), ensuring that any identified effects were not due to background neural dynamics. Second, the model included a matrix ($T_{auto}$) that models self-transitions to control for auto-correlation (**Fig. S3b**). The evidence for all sequences of interest was then quantified by:

$$\boldsymbol{B} = \sum_r Z(r) * T_r \quad (13)$$

where $r$ is the number of all regressors included in the second stage, as specified above. $Z$ is the scalar regression coefficient quantifying the evidence that the hypothesized transitions, $T_r$ predict the empirical transitions, $\boldsymbol{B}$ (i.e. sequenceness strength). Note, this estimate of sequence strength is a relative quantity. An estimate of zero for state *i* to state *j*, for example, does not mean there is no replay of $i \rightarrow j$, it means there is no stronger replay of $i \rightarrow j$, than that of other transitions. Repeating the regression of Equation 13 at each time lag ($\Delta t$ = 10, 20, 30, ... , 600 ms) results in timecourses of both forward and backward sequence strength as a function of time lag, where smaller lags indicate greater time-compression of replay (**Fig. S3b**). For each trial, sequenceness results were z-scored across lags; note that this had no qualitative effect on the learning phase results. A subsequent model substituted the full transition matrices with matrices that separated the paths from the two worlds (current, other) in the forward ($T_{Fcurrent}$, $T_{Fother}$) and backward ($T_{Bcurrent}$, $T_{Bother}$) direction and was otherwise identical. Subsequently, in the initial lag localization method (below), sequenceness was averaged across trials. All other analyses utilized per-trial sequenceness values.

Unlike rodent electrophysiology, we cannot detect discrete replay events given that we are not measuring spiking data, but an LFP-like continuous signal. This is another reason we utilize linear modelling to assess the sequence strength on average across a trial period. Whilst the sequenceness method was originally tested on longer time periods > 10 s (Kurth-Nelson et al., 2016; Liu et al., 2019), recent work has demonstrated its applicability to shorter time periods (Wimmer et al., 2020; Liu et al., 2021b), where simulations have verified an ability of the method to discriminate between true sequences and noise. As our analysis used short trial periods and not long rest periods, we did not find a strong alpha rhythm and did not control for alpha in the sequenceness analyses (Liu et al., 2019; Wimmer et al., 2020). Note that for analyses of relatively short state sequences, as found in our task, sequenceness values could be driven selectively by preferential replay of early or late transitions, but we did not find that this was the case.

To localize forward and backward time lags of interest for the primary trial-by-trial regression analyses, we identified significant lags using non-parametric permutation tests with shuffled transition matrices (n = 100). Shuffled transition matrices were randomly generated for each trial to include only invalid transitions but to otherwise follow the structure of the true matrix. To match the structure of the true paths, each shuffled transition matrix was required to have four independent paths. Thus, each shuffled matrix contained eight entries (two per path). This ensured that each stimulus would appear in a sequence a maximum of once per permutation and that true sequential triplets of states were formed across states.

For each permutation, sequenceness values were averaged across trials (within-participant), and then across participants. Then, significance thresholds were determined in two steps (separately for forward and backward sequenceness). First, for each permutation, we computed the maximum permutation-derived sequenceness value across all time lags. Second, across the resulting set of 100 maximum values, we applied a 95% threshold to determine the corrected significance level. By using the maximum value across lags in the first step, the resulting threshold provides for a statistical correction across all lags at the 0.05 level (Liu et al., 2021a). This approach has been validated in both simulation and empirical data (Liu et al., 2019; Wimmer et al., 2020; Liu et al., 2021a). As human studies of sequential state reactivation have only found evidence for replay at relatively short lags (e.g. Kurth-Nelson et al., 2016; Liu et al., 2019), a finding which we replicate here, we display results up to a lag of 350 ms. Further, given the above rationale, in post-hoc analyses for lags not identified in the lag localization step, we use this shorter range of lags (10 – 350 ms) as the basis for Bonferroni multiple comparisons correction.

**Identifying Replay Onsets**

For follow-up source localization and time-frequency analyses, replay onsets were defined as moments when a strong reactivation of a stimulus was followed by a strong reactivation of the next (or preceding) stimulus in the sequence from a path (Liu et al., 2019; Wimmer et al., 2020). As described in the Results, a different direction and lag was the focus of the planning and reward feedback period analyses. We identified the stimulus-to-stimulus time lag $\Delta t$ at which there was maximum evidence for sequenceness (as described above). We first generated a matrix Orig as

$$Orig = X \times T \tag{14}$$

where X is the [time x state] reactivation matrix, and T is the task transition matrix. The transition matrix T defines the mapping between the task state corresponding to column i in X, and column i in Orig (specifically, column i in Orig is the reactivation timecourse of the state that 'precedes' state i in T). We then shifted each column of X by the relevant replay lag, Δt, to generate another matrix Proj,

$$Proj = X(\Delta t) \tag{15}$$

where row i of Proj corresponds to row i+[lag] ms of X. Multiplying Proj and Orig elementwise, and summing over the columns of the resulting matrix, therefore yields a [time x 1] vector, R, where each element, t, corresponds to the evidence for a two-state replay with given lag, starting from any task state at time t.

$$R_t = \sum_i^s Orig_{ti} * \ Proj_{ti} \tag{16}$$

We then identified putative replay onsets. Within-participants, we thresholded *R* at its $95^{th}$ percentile to only include high-magnitude putative replay onset events. We also imposed the constraint that a replay onset event must be preceded by a 200 ms pre-onset baseline period exhibiting summed reactivation evidence < $90^{th}$ percentile at each time point.

**MEG Source Reconstruction**

Source reconstruction (beamforming) was performed in SPM12 and FieldTrip utilizing OAT. Forward models were generated on the basis of a single shell using superposition of basis functions that approximately corresponded to the plane tangential to the MEG sensor array.

Linearly constrained minimum variance beamforming (Van Veen et al., 1997) was used to reconstruct the epoched MEG data to a grid in MNI space, sampled with a grid step of 5 mm. The sensor covariance matrix for beamforming was estimated using the preprocessed data in broadband power across all frequencies in the lower range 1.5–47.3 Hz.

For the replay onsets analysis, we computed baseline activity as the mean power averaged over -100 ms to -50 ms relative to replay onset. All non-artifactual trials were baseline corrected at source level. We looked at the main effect of the initialization of replay. This analysis was conducted separately to investigate forward replay events in the planning period and backward replay events in the feedback period. To increase power, primary analyses examined putative replay onset events across both worlds, while secondary analyses examined current world replay onsets during planning and other world replay onsets during feedback.

The statistical significance of clusters identified in the beamforming analysis was calculated using non-parametric permutation tests in OSL to identify clusters significant at $P_{FWE} < 0.05$ (whole-brain corrected; cluster-defining threshold $p < 0.01$ ($t = 2.81$); 5000 permutations). We expect power increases in source space to reflect increases in underlying neural activity. Replay onsets are defined by high evidence for internal generation of a visual stimulus. As expected, previous reports find robust replay-onset-related activity in the visual cortex in addition to the MTL (Wimmer et al., 2020; Liu et al., 2021b). Further, this internally-generated activity overlaps with that detected for actual stimulus presentation (Wimmer et al., 2020) and generally corresponds to regions showing increased visual-evoked signal in fMRI and electrophysiological studies. Finally, we assume that continuous regions of activity extending anterior from the visual cortex to the medial temporal lobe would reflect the same positive direction of underlying neural activity.

**Time-frequency analyses**

A frequency decomposition (wavelet transformation) was computed separately for the planning and reward feedback period in every trial. From this data, we extracted power changes surrounding the putative replay onset events (± 200 ms) identified using the preceding method. To increase power, primary analyses examined putative replay onset events across both worlds, while secondary analyses examined current world replay onsets during planning and other world replay onsets during feedback. We examined evidence across participants for (mean) power increase at replay onset compared to a pre-onset baseline (from –200 to –150ms before replay onset) within a frequency range of interest of 120-150 Hz, in the ripple band that has been previously associated with replay events (Liu et al., 2019; Liu et al., 2021b; Nour et al., 2021) as well as a theta band range of interest (5-8 Hz). Within these bands of interest, we tested individual difference correlations between replay onset (± 2.5ms) and individual differences in the replay-behaviour relationships previously identified. A cluster-based permutation analysis implemented in OSL separately tested whether there were significant clusters associated with replay onset across the full time-frequency range. The clustering algorithm used an initial threshold of $p < 0.01$ (corresponding to a t-threshold of 2.81) and an alpha level of 0.05.

In control analyses to test for the specificity of our results to sequential replay, we used the above trial-wise frequency decomposition for the planning and reward feedback periods. Within each participant, at each timepoint and frequency, we estimated the correlation across trials with the planning and feedback variables of interest: at planning, the benefit of generalization and state value; at feedback, rarity of other world experience. These results were submitted to cluster permutation analyses as in the preceding replay onset time-frequency analyses.

**Non-sequential reactivation analyses**

In two additional control analyses, we tested for the specificity of our results to sequential replay. We used the same time by state classifier evidence for the planning and reward feedback periods that also served as input for the sequenceness analyses. The first set of control analyses examined mean classifier activity across the time period, separately for the current and other world, averaging across the states within a path. Additional checks examined each of the three path states separately. As in the sequenceness analysis, these measures reflect the relationship across the full time period of interest (planning, feedback) for each trial. The second set of control analyses examined pair-wise 'clustered' reactivation of states present on the same path, following previous work (Wimmer et al., 2020). Clustered reactivation was calculated for each trial as the zero-lag correlation of classifier evidence between two elements of a given path. Correlation values were z-transformed and then these z-values were averaged across the potential pairs in a path or world. Both of these trial-by-trial control measures were analyzed using multilevel models

In a more temporally fine-grained analysis, we also examined the relationship between reactivation at each time point (in 10 ms resolution) and variables of interest. For statistical comparison, these results were submitted to cluster permutation analyses as used in the preceding replay onset time-frequency analyses.

**Representational similarity**

We used Representational Similarity Analysis (RSA) (Diedrichsen and Kriegeskorte, 2017) to investigate the emergence of a representation of task structure from the stimulus localizer (pre-learning) to reward learning, following previous procedure (Nour et al., 2021). We utilized this analysis because we lacked a post-task localizer and because RSA may be more sensitive to effects due to abstract structure than sequenceness analyses in short task periods (versus longer rest durations in previous studies). First, for each trial in each session we subtracted the mean of the 100ms preceding stimulus onset from the pre-processed MEG data (baseline correction); for each of the three stimuli presented in reward learning paths, this correction was performed per-stimulus. For each sensor, we then z-scored over all trials and time points relative to stimulus-onset (t, -100 to +1500 ms). We then regressed the [trial x 1] neural data, $Y(s)_t$ (from time point, t, and sensor, s) onto a session design matrix, X, denoting the stimulus label of each trial (dummy coded) (Luyckx et al., 2019),

$$Y(s)_t = X * \beta(s)_t \qquad (17)$$

and used the resulting [stimulus x 1] vector of regression weights, $\beta(s)_t$, as an estimate of the unique activation for each stimulus, in sensor s at time point t. Repeating this procedure over all sensors yielded a [sensor x stimulus] matrix at each time point. We then calculated the Pearson correlation distance between the sensor patterns for each pair of stimuli (columns). This generated a symmetrical [12 x 12] Representational Dissimilarity Matrix (RDM) at each time point (Deuker et al., 2016). We conducted this procedure identically in both stimulus localizer (SL) and reward learning (R), enabling us to calculate the learning-induced change in representational similarity (similarity change) at each time point, $\Delta S_t$, as

$$\Delta S_t = RDM(SL)_t - RDM(R)_t \quad (18)$$

where entry $s_{ij}$ of $\Delta S_t$ quantifies the post-learning similarity increase between evoked signals for stimuli i and j, at time t (Deuker et al., 2016). $\Delta S_t$ was then smoothed over time with a 50 ms Gaussian kernel (Luyckx et al., 2019).

Finally, we used a second multiple regression to quantify the variance in $\Delta S_t$ that was uniquely explained by an abstracted representation of ordinal position for each participant at each time point. The position predictor was separated into two matrices, capturing position similarity across worlds and position similarity within the same world. Path identity was also included in the model. The multiple regression thus included 3 predictor variables corresponding to each of the hypothesized representational patterns, plus an intercept (see **Fig. S9**). Note that motor response to the stimuli was not a confound; participants responded with a button press whether stimuli were presented on the left or right side of the screen, but the left/right position was assigned randomly per trial per stimulus.

We used nonparametric tests to identify time windows (clusters) with significant positive evidence for each predictor, correcting for multiple comparisons over time (Nour et al., 2021). Specifically, for a given predictor (e.g., position), at each time point post-stimulus onset we conducted a separate one-sample t test on the representational effect (regression weights) over participants, to obtain the evidence for an effect > 0 (i.e., t-value). We computed the sum of t-values within each continuous stretch of time points exhibiting a positive effect at $p < 0.05$. We then repeated this procedure for 1000 permutations, on each occasion applying the same novel shuffling to the rows and columns (stimulus labels) of $\Delta S_t$ prior to the second regression. The shuffled order was consistent across time within each permutation to preserve temporal smoothness in visually evoked neural data. We then extracted the maximal sum-of-t value for the group mean effect in each permutation (matching the procedure applied to the unpermuted data), to generate an empirical null distribution for the predictor in question. We expected any position representation effect to be positive. A suprathreshold positive cluster in the unpermuted data was deemed significant at $P_{FWE} < 0.05$ if its sum-of-t values exceeded the $95^{th}$ percentile of this empirical null distribution (Eldar et al., 2018; Nour et al., 2021). We had no expectations about the direction of any path identity representation effect, so clusters were judged to be significant at $P_{FWE} < 0.05$, two-tailed, if its sum-of-t values exceeded 97.5% (or fell below 2.5%) of this null distribution.

**Multilevel regression models**

Multilevel models relating sequenceness to trial-by-trial variables of interest were estimated using R, as further detailed above for the behavioral models. In these models, the independent variables were the relevant and control sequenceness strength values, e.g. current and other world paths sequenceness. All relevant sequenceness measures were included in the same model. To allow for valid comparisons, where analyses compared regression coefficients or computed interactions, the relevant variables were z-scored across trials.

**Statistical correction and null effects**

For permutation-based significance measures of mean overall replay, see the relevant section above. For trial-by-trial analyses, multilevel models were implemented in R, as described above. Other tests of means or fit parameters utilized t tests except where the noted. All reported tests are two-tailed unless otherwise noted. One-tailed tests were used when we had an *a priori* expected direction of a comparison, including modulation of current versus other world replay or planning versus feedback period replay. For the exploratory analyses across additional replay time lags, significance was determined using Bonferroni correction for the number of comparisons (34, where calculated lags ranged from 10 to 350 ms). Contrasts of multilevel model regression coefficients were estimated using the glht function in the multcomp package; note that all variables entered into regressions were z-scored to allow for valid contrasts. For comparisons of softmax inverse temperature parameters, as expected, the distribution of the inverse temperature parameter was not normal, so we used Wilcoxon signed rank tests. For results of interest, we additionally tested whether non-significant results were weaker than a moderate effect size using the Two One-Sided Test (TOST) procedure (Schuirmann, 1987; Lakens, 2017), as implemented in the TOSTER library in R (Lakens, 2017). We used bounds of Cohen's d = 0.60, where power to detect a medium- or larger-sized effect in the group of 24 participants was estimated to be 80%.

### Supporting text

#### Choice regression analysis

We used two behavioral regressions approaches. The first, discussed in the Results, predicted stay decisions based on preceding reward. The second regression that we describe here was similar, but instead predicted the identity of the option selected based on preceding reward (Doll et al., 2015). As expected, we found a strong overall effect of previous reward receipt on choice (reward effect on choice, multilevel logistic regression coefficient $\beta = 0.529$ [0.420 0.637]; $z = 9.582$, $p < 0.001$). We then tested for any model-free influence via an interaction between reward and start state change. A model-free influence would be evident if the effect of previous reward on choice is be greater for choices where the current start state matched the previous start state. We found no such difference (reward effect and same start state interaction $\beta = 0.036$ [-0.0248 0.0961]; $z = 1.156$, $p = 0.248$; TOST $p = 0.44$). We also extracted individual participant interaction coefficients, which serve as an index of model-based behavior (Doll et al., 2015). As expected, this derived model-based index was significantly correlated with the model-based weighting parameter $w$ from the RL model ($r = 0.663$, $p = 0.0004$).

#### Behavioral effect of trial-to-trial world change

We expected that when trials changed from one world to the other, choice would be negatively affected, as a delay between repetitions of the same world would be expected to decrease the ability of working memory to support behavior (Collins and Frank, 2012; Wimmer et al., 2018). Indeed, alternating from one world to another had a negative effect on performance, decreasing the influence of previous reward on choice ($\beta = -0.0482$ [-0.0883 -0.0081]; $z = -2.355$, $p = 0.019$). We found a numerically stronger negative effect of world alternation when we accounted for the number of intervening alternate world trials ($\beta = -0.0881$ [-0.0144 0.1898]; $z = -3.952$, $p = 0.00008$).

#### Behavioral effect of cued stakes

We asked whether reward stakes cued at the time of choice increased model-based behavior (**Fig. 1b**), following previous research (Kool et al., 2017). Unfortunately, we found that the pseudo-randomization of stakes and reward point drift across trials led to an unintended correlation between model-predicted option values and stakes (best fit model-based model-derived choice value, $p = 0.0066$; difference in values, $p = 0.0012$). As this relationship between signaled stakes and internally-estimated values may have influenced participants' choice behavior in unpredictable ways, we are not confident in drawing conclusions from the stakes manipulation. For completeness, and with this confound in mind, we present basic behavioral analyses here.

We found a non-significant positive influence of stakes on choice, such that high stakes tended to increase the influence of previous reward on choice (interaction $\beta = 0.0391$ [-0.0066 0.0849]; $z = 1.676$, $p = 0.0937$). We tested for but did not find an across-participant relationship between this influence of stakes and our indices of model-based behavior (start state interaction correlation $r = 0.217$, $p = 0.308$; *w* parameter correlation $r = 0.153$, $p = 0.477$). Within-participant, stakes was also not correlated with reaction time ($p = 0.745$). We also fit a hybrid RL model with two model-based weighting parameters *w*, one for high and one for low stakes trials, based on previous work showing a positive influence of stakes on the model-based weighting parameter (Kool et al., 2017). We found no impact of stakes on *w* ($p > 0.47$). Further, we found that current trial stakes did not modulate learning: in a model with a separate learning rate for high stakes and low stakes trials, we found no difference in learning rate and no increase in model fit to behavior.

**Model-based behavior and weighting parameter**

The deterministic structure in our task, among other features, sets apart this design from the probabilistic alternative by actually incentivizing model-based behavior (Kool et al., 2016; Kool et al., 2017; Patzelt et al., 2019). Indeed, we found in simulations that model-based behavior led to higher reward earnings (**Fig. S2b**).

Beyond the regression and RL model evidence presented in the Results, additional support for the strength of model-based behavior was evident in the hybrid model where the model-based weighting parameter *w* was skewed toward 1 (median =

0.802; **Table S1**). Notably, this degree of model-based weighting is higher than reported in similar versions of this paradigm (Doll et al., 2015; Kool et al., 2016), as well as versions with probabilistic state transitions (Daw et al., 2011). This indicates that participants' behavior in the present task may be more model-based than in preceding studies, potentially reflecting a deterministic task structure and multi-day training (Liu et al., 2021b). However, even with low variability in the model-based weighting parameter *w*, we found that *w* positively correlated with mean total reward earnings ($r = 0.826$, $p < 0.0001$), suggesting that our task design effectively incentivizes model-based behavior (Kool et al., 2016).

**No change in model-based behavior across time**

We found a consistent model-based effect across time (first half β = 0.467 [0.320 0.613]; z = 6.255, p < 0.0001; second half β = 0.515 [0.376 0.654]; z = 7.238, p < 0.0001; trial interaction β = 0.059 [-0.089 0.207]; z = 0.783, p = 0.434; TOST p = 0.021). Thus, for choices requiring generalization, the influence of previous reward on subsequent choice if anything numerically increased over time. However, we found that the strength of a model-free influence, measured on choices not requiring generalization, significantly increased across time (first half β = 0.448 [0.316 0.580]; z = 6.675, p < 0.0001; second half β = 0.813 [0.598 1.029]; z = 7.403, p < 0.0001; trial interaction β = 0.234 [0.022 0.447]; z = 2.160, p = 0.031), suggesting participants were better at using previous feedback to influence same-state choices.

In reinforcement learning models, we found no change in the fit of the model-based model across the task (first half mean likelihood = 32.8; second half = 33.4; p = 0.691; TOST p = 0.009) and also no change in the fit of the model-free model across the task (first half = 38.4; second half = 40.4; p = 0.313; TOST p = 0.034). In the hybrid model, the model-based weighting parameter *w* also did not change (first half median = 0.878; second half = 0.842; difference p = 0.249; TOST p = 0.046). Thus, participants' signature of model-based learning, reflecting structure knowledge which allows for the generalization of reward when starting in a different state, did not change. For model-free learning, one of two measures indicated an increase over time, indicating an increased ability to use direct reward feedback to guide subsequent choice in the same start state.

**MEG: No overall planning sequenceness during initial structure learning**

We examined sequenceness during participants' initial exploration of the two worlds preceding the reward learning phase. In this phase, participants received no reward feedback and their only goal was to learn the state-to-state transitions between the objects. For results of analyses of replay during the inter-trial interval, see the main **Results** section. During the planning period we found no above-threshold sequenceness. We found no significant effects when restricting analyses to the second half, and numerically positive but non-significant increases when testing the difference between the second and first half.

**Across-sequenceness correlations**

In the learning phase, we examined whether the different replay measures (planning and feedback forward and backward replay) were correlated with each other within-participants at a trial-by-trial level. During planning, for forward sequenceness with a 70 ms state-to-state lag, we found no significant correlation between current and other world sequenceness ($p = 0.435$) or between chosen and non-chosen path forward sequenceness ($p = 0.183$).

At feedback, for backward sequenceness with a 40 ms state-to-state lag, we found no significant within-participant replay correlation between the current and other world path sequenceness ($p = 0.710$) or between chosen and non-chosen path sequenceness ($p = 0.530$). For forward sequenceness with a 70 ms state-to-state lag, we also found no significant correlation between current and other world sequenceness ($p = 0.343$) or between chosen and non-chosen path sequenceness ($p = 0.299$).

Across periods, we tested for within-participant relationships between replay during planning and replay after feedback at a trial-to-trial level for current and other world sequenceness. We found no significant correlations between forward sequenceness measures, backward sequenceness measures, or across forward and backward sequenceness measures (p-values > 0.132). Further, we found no overall relationship between previous other world feedback backward replay and current world forward replay (overall, or selective to world change trials). These results indicate that replay measures were not significantly related by simple positive or negative correlations, and that there was no significant evidence of a trial-by-trial relationship between replay strength at planning and at feedback.

Finally, we examined across-participant (individual difference) relationships between mean sequenceness measures. During planning, mean forward replay at a 70 ms lag was not significantly correlated with reverse replay at a 40 ms lag (measured across all paths in the current and other world; r = 0.306, p = 0.147). During feedback, mean forward replay at a 70 ms lag was not significantly correlated with reverse replay at a 40 ms lag (r = -0.245, p = 0.249). Across periods, we found that mean forward replay during planning was correlated with the same measure at feedback (70 ms lag; r = 0.694, p = 0.00017, uncorrected for multiple comparisons), potentially reflecting the finding of significant forward replay in both periods. Mean backward replay was not correlated across periods (40 ms lag; r = 0.0169, p = 0.938).

**No relationship between planning period replay and choice identity**

We also explored whether relative sequence strength for the to-be-chosen versus non-chosen paths related to subsequent path choice. We found no evidence that sequenceness was predictive of choice overall, or for choices in different types of trials (stay, switch, or exploring low-value choices). Thus, overall sequenceness strength was similar for chosen and non-chosen paths (comparison -0.0084 [-0.0612 0.0445]; $t_{(23)} = -0.327$, p = 0.746; TOST p = 0.0078). We also found no relationship between mean path state reactivation (either across the whole path, or examining each state separately) and choice (**Fig. S3b**), or between non-sequential paired reactivation and choice.

We speculate that it may be difficult to find general relationships between replay and option selection, especially in commonly-employed two-alternative choice paradigms. In these tasks, path evaluation and a correlated replay signal could positively bias choice because of expectation of relatively high reward from a given option (Wikenheiser and Redish, 2015), or could negatively bias choice because of expectation of relatively low reward from the other option (Wu et al., 2017), with these effects potentially varying across participants and trials.

### Control analyses demonstrating specificity of replay effects

**Model-based generalization and replay.** We tested whether individual differences in model-based decision making (indexed by the interaction with start state and reward) were related to an increase in sequenceness when generalization was more beneficial. We found no across-participant relationship between individual differences in model-based behavior and the increase in current world replay by generalization ($p = 0.873$). However, in an exploratory analysis we found that individual differences in model-based behavior were associated with a decrease in other world replay by generalization ($r = 0.524$, $p = 0.0085$). We also asked whether behavioral variability in the form of reaction time slowing for generalization (start state change) trials related to forward sequenceness. A decreased reaction time cost of start state change was numerically, but not significantly, related to stronger current world sequenceness (interaction $\beta = -0.3956$ [-0.7989 0.0078]; $z = -1.922$, $p = 0.0546$).

Control analyses for the replay–generalization effect verified that the relationship between replay and a benefit for model-based planning was specific to forward replay with a 70 ms state-to-state lag during the planning period (see **Table S3** for a summary). First, during planning, current world backward replay at a 40 ms lag showed no relationship to model-based generalization benefit ($p = 0.729$; TOST $p = 0.008$), and the forward effect was significantly stronger ($z = 2.267$, $p = 0.012$, one-tailed). During planning, current world backward replay at a 70 ms lag showed no relationship to generalization benefit ($p = 0.406$; TOST $p = 0.023$), and the forward replay effect was significantly stronger than the backward effect ($z = 3.175$, $p = 0.0008$, one-tailed).

Second, at feedback, we found no significant relationship between backward replay at a 40 ms state-to-state lag and generalization benefit ($p = 0.119$), and the planning period effect was significantly stronger than this feedback effect (z = 2.891,3 p = 0.002, one-tailed). We also found no relationship between feedback period forward replay at a 70 ms state-to-state lag and generalization benefit (p = 0.537, TOST p = 0.015; current world planning versus feedback forward replay, z = 1.192, p = 0.117, one-tailed).

Third, we found no relationship between control non-sequential reactivation measures and the relative benefit of model-based generalization. Mean planning period activity strength for current world path states was not related to generalization (p = 0.240; TOST p = 0.046; forward replay versus reactivation difference z = 01.532, p = 0.063, one-tailed). Additionally, non-sequential “paired” reactivation of path states in the current world was also not related to generalization (p = 0.547; TOST p = 0.0142; forward replay versus reactivation difference z = 1.943, p = 0.026, one-tailed). In a separate analysis of time-frequency data, we also found no relationship with model-based generalization (n.s. using permutation-based cluster correction).

Fourth, in exploratory analyses, we also investigated planning period backward replay at a 160ms state-to-state lag. In a recent publication, several of the authors of the current paper found that backward replay after feedback was related to learning (Liu et al., 2021b). As discussed in the main text, the current paradigm was specifically designed to better elicit planning-related replay prior to choice. We found no relationship between 160 ms lag backward replay and the benefit of model-based generalization (p = 0.195, TOST p = 0.057) and the forward 70 ms relationship was significantly stronger than the 160 ms lag backward effect (difference z = 1.850, p = 0.0321, one-tailed). Forward replay at a 160ms state-to-state lag was also not related to model-based generalization (p = 0.314; TOST p = 0.033) and the forward 70 ms relationship was significantly stronger than the 60 ms lag forward effect (difference z = 3.467, p = 0.0026, one-tailed).

Finally, we compared the predicted effects due to generalization with an alternative account related to memory retrieval and maintenance (‘refreshing’). This alternative uses the rarity (inverse recency) of a given start state, analogous to the world rarity variable linked to feedback replay. A planning account of our task implementation predicts that planning should be driven by the binary effect of start state change versus no change. However, an alternative account based on memory retrieval and maintenance predicts that current world replay should relate to the overall frequency of experiencing a given start state. Here, replay would reflect the fact that retrieval and maintenance demands are higher for states that have less experience in the recent past. The memory retrieval model thus predicts that replay would be related to a graded variable tracking rarity of experience, instead of the binary effect of start state change (generalization) predicted by the planning account.

Note that in our task, reward points drifted relatively rapidly across trials. As a consequence, the relevance of previous experience in a given start state for estimating current values decays relatively quickly when alternative start state trials intervene. Thus, when the start state changes, beyond reacting to the immediate demands for generalizing, there is little to no advantage in adjusting replay-related planning based on how frequently a start state has been experienced.

We constructed a control 'recency of experience' learning model which tracks start shape frequency and updates this variable on a trial-to-trial basis according to a learning rate parameter. We then tested whether recency is more strongly related to planning period replay. First, we looked at a weaker version of the memory model, tracking recency of experience only within a given world. The planning account is similar to this model with a recency learning rate set to 1. As the recency learning rate decreases, the recency variable is less and less correlated with the binary start state change variable. If the memory account was a better explanation of the planning replay effect, the peak effect would occur with a learning rate below 1.0. However, as the recency learning rate decreased, the relationship to sequenceness systematically and linearly decreased and was no longer significant when the learning rate was below 0.20. Thus, this pattern of results supports the planning account over a memory retrieval account.

We then tested a stronger version of the alternative memory retrieval and maintenance demand model: whether replay tracked the actual recency of start state experience across trials, accounting for intervening experience in the alternative world. In this model, in addition to adjustments due to intervening different start state trials within a world, the rarity of both start states in a world increases during the presentation of other world trials. The memory account would predict a peak in the relationship at some learning rate below 1.0, but this was not the case. We found that the relationship with sequenceness was numerically weaker than the within-world recency control model, above, and was also no longer significant when the learning rate was below 0.20. Finally, for a control model with no free parameters, we tested an alternative model that predicts an increase of current world replay when the world changes from trial to trial. The relationship between current world replay and world change was equivalent to zero ($p = 0.341$; TOST equivalence $p = 0.0294$). These patterns of results support the planning account over an alternative memory retrieval account.

**Option value and replay.** Control analyses verified that the relationship between replay and value was specific to forward replay during the planning period (see **Table S3** for a summary). Further, separating the paths and the value measure, we found that within current world paths, chosen path sequenceness was correlated with the estimated value of the chosen option ($\beta = 0.0366$ [0.0008 0.0724]; $t = 2.00$, $p = 0.0452$), while non-chosen path sequenceness was not correlated with the estimated value of the non-chosen option ($\beta = 0.0210$ [-0.0145 0.0565]; $t = 0.876$, $p = 0.247$; TOST $p = 0.044$). We found no significant relationship between current world replay and the difference in option values ($\beta = 0.0251$ [-0.0131 0.0632]; $t = 1.289$, $p = 0.198$; TOST $p = 0.056$).

We also found no relationship between the control non-sequential reactivation measures and the benefit of model-based generalization. Mean planning period activity strength for current world path states was not related to state value ($p = 0.686$; TOST $p = 0.009$; replay versus reactivation difference $z = 0.999$ , $p = 0.159$, one-tailed). Additionally, non-sequential “paired” reactivation of path states in the current world was also not related to state value ($p = 0.686$; TOST $p = 0.009$; replay versus reactivation difference $z = 1.136$, $p = 0.128$, one-tailed). In a separate analysis of time-frequency data we also found no relationship to state value (n.s. using permutation-based cluster correction).

**Recent experience (memory preservation) and replay.** Control analyses verified that the relationship between backward sequenceness and the rarity of recent experience was specific to backward replay in the feedback period (see **Table S3** for a summary). First, during the planning period, we found no modulation of other world forward replay at a 70 ms state-to-state lag and recent experience (p = 0.988; TOST p = 0.004), while the feedback effect was significantly stronger than the planning effect (z = 1.778, p = 0.0378, one-tailed). We also found no evidence for modulation of planning period other world backward replay at a 40 ms lag (p = 0.606; TOST p = 0.012), while the feedback effect was significantly stronger than the planning effect (z = 1.972, p = 0.024, one-tailed).

Second, during the feedback period, we found no relationship between other world forward replay at a 70 lag and rarity (p = 0.690; TOST p = 0.0092; current world p = 0.857; TOST p = 0.0056), while the backward replay effect was non-significantly stronger than the forward effect (z = 1.50, p = 0.067, one-tailed). Separately, for the backward replay effect of interest, we found no interaction between the rarity effect of interest and other world value (p = 0.778; TOST p = 0.007). This indicates that a modulation of replay by rarity is unrelated to (low) value, a potential confound in previous related reports in rodents (Gupta et al., 2010; Carey et al., 2019).

Third, we found no relationship between the control non-sequential reactivation measures and the rarity of recent experience. Mean planning period activity strength for current world path states was not related to recent experience (p = 0.305; TOST p = 0.034; difference z = 0.969, p = 0.167, one-tailed). Additionally, non-sequential "paired" reactivation of path states in the current world was also not related to recent experience (p = 0.850; TOST p = 0.006; difference z = 1.539, p = 0.124, one-tailed). In a separate analysis of time-frequency data, we also found no relationship with rarity of experience (n.s. using permutation-based cluster correction).

Fourth, in exploratory analyses, we also investigated feedback period backward replay at a 160ms state-to-state lag, as identified recently at feedback and as described above (Liu et al., 2021b). We found no relationship between 160 ms lag backward replay and the rarity of recent other world experience (p = 0.438, TOST p = 0.021), while the 40 ms lag backward effect of interest was numerically stronger than the control 160 ms lag effect (z = 1.208, p = 0.114, one-tailed).

**Reinforcement learning effects of memory preservation signal**

An RL model incorporating neural signals examined the link between feedback period other world replay and memory preservation by allowing replay to modulate next choice in that world. We predicted that stronger preceding replay of the other world would translate to a higher softmax inverse temperature, reflecting lower uncertainty associated with structure memory and learned values. As expected, the distribution of the inverse temperature parameter was not normal (given constraints in fitting of [0 Inf]), so we tested our hypothesis using the Wilcoxon signed rank test (one-tailed, reflecting our a priori prediction). As described in the Results, when the world changed from trial to trial, we found a significantly higher inverse temperature on trials with strong preceding replay versus not. Control analyses verified that the influence of preceding feedback replay on softmax inverse temperature (uncertainty) in the RL model was specific to backward replay in the feedback period.

We conducted several control analyses using an alternative model or substituting alternative replay measures to determine the specificity of this finding. First, we tested whether the replay modulation of inverse temperature was selective to world-change trials as expected. In this model, instead of world-change trials, the additional inverse temperature parameters applied to world-same trials with stronger preceding other world replay and with weaker preceding replay. We found no effect of replay modulating uncertainty (via inverse temperature) on world-same trials ($p = 0.753$, one-tailed, Wilcoxon signed rank test).

Second, we examined the effect of different replay signals at feedback. The presence of previous trial current world replay – reflecting the "other world" structure on the current choice – did not relate to an increase in the inverse temperature parameter ($p = 0.173$, one-tailed). Further, forward, instead of backward, previous trial replay of the relevant other world also did not relate to a significant increase in the inverse temperature parameter (40 ms lag; $p = 0.410$, one-tailed). These analyses indicate that the feedback replay effect in the RL model was specific to sequential reactivation of states in the relevant world in the expected backward direction.

Third, we examined the effect of planning period backward replay of the current world, which has the same content as previous trial "other world" replay. As expected, we found no significant increase in the inverse temperature parameter (40 ms lag, $p = 0.274$, one-tailed). We also found no effect of planning period forward replay of the current world on the inverse temperature parameter (40 ms lag; $p = 0.421$, one-tailed; 70 ms lag; $p = 0.443$, one-tailed). These analyses indicate that the feedback replay effect in the RL model was specific to the feedback period. Moreover, this selectivity to the feedback period supports the proposal that the relatively low computational demands after feedback receipt favor a memory preservation signal.

Finally, as reported in the Results, we examined whether this effect on the inverse temperature parameter was specific to replay, or a general function of rare experience of a world. It is possible that aspects of decision making could differ for worlds that have not been experienced recently, and that this could give rise to changes in the inverse temperature parameter independent of any preceding replay differences. Analogous to the replay measure, we thresholded rarity at the $60^{th}$ percentile to create a binary variable and examined any influence on the inverse temperature on choices when returning to the other world. We found no difference in inverse temperature due to more versus less rare choices ($p = 0.246$, one-tailed).

**Feedback period reward response and forward replay**

In the feedback period, we also identified significant forward replay at a 70 ms lag. We examined whether current or other world forward replay after feedback was modulated by reward and other variables of interest (**Table S3**). We found a non-significant but numerically negative relationship between reward feedback receipt and current world forward sequenceness ($\beta = -0.0875$ [-0.2045 0.0295]; $t = -1.467$, $p = 0.143$; **Table S3**). This effect was similar for chosen path and non-chosen path sequenceness (chosen path $p = 0.213$; non-chosen path $p = 0.287$). There was no significant relationship between reward and other world sequenceness ($\beta = 0.0641$ [-0.0596 0.1878]; $t = 1.016$, $p = 0.310$). Forward sequenceness for current or other world paths also did not significantly relate to model-derived reward prediction error (p-values > 0.262). Additionally, for backward replay with a 40 ms lag we found no effects of reward prediction error (p-values > 0.611; **Table S3**). We also examined the replay signal in the initial period of feedback processing (the first 2.5 s, excluding the initial 160 ms to allow for visual processing). Here, all relationships with reward and reward prediction error were equivalent to the null (via the TOST procedure).

Next, we examined the effect of surprising outcomes, using unsigned (absolute value) reward prediction error. We found that other world forward sequenceness was negatively related to unsigned reward prediction error (β = -0.0077 [-0.0348 0.0490]; t = -2.822, p = 0.0048), with no effect on current world sequenceness (β = 0.0071 [-0.1020 -0.0184]; t = 0.331, p = 0.741; difference z = 2.251, p = 0.0122, one-tailed). These results may suggest that when feedback surprise in the current world is high, relative current versus other world replay is higher, which may potentially aid processing of surprising outcomes. Speculatively, it is possible that a relative decrease in other world versus current world replay after surprising feedback may assist a planning or planning preparation function. However, our task was designed to promote planning when options are presented, in contrast to related recent work on feedback processing (Liu et al., 2021b); thus, measurable modulations of current world replay may be more likely to be found in the planning period.

**Feedback period backward replay with a 160 ms lag, as in Liu et al. 2021**

In a recent publication, several of the authors of the current paper found that backward replay after feedback was related to learning (Liu et al., 2021b). In the current data, in the feedback period we did not find significant backward (or forward) replay at or near a 160 ms lag. As described above, we found no link between 160 ms lag replay and either the benefit of model-based generalization or the rarity of recent other world experience.

When further examining 160 ms lag feedback backward replay, we found positive but non-significant effects for reward for both current and other world paths (160ms lag backward replay, current world, t = 1.6469, p = 0.0997; other world, t = 1.7835, p = 0.0746). We found no significant relationship with reward prediction error, although there was a numerically positive effect (160ms lag backward replay, current world, t = 1.704, p = 0.089; other world, t = 0.817, p = 0.414). We also found no significant correlation with reward prediction error when separately looking at replay for the chosen and non-chosen paths in the current world (chosen p = 0.084; non-chosen p = 0.392). We found no relationship with the absolute value of reward prediction error. We also examined this signal in the initial period of feedback processing (the first 2.5 s, excluding the initial 160 ms to allow for visual processing). Here, we found numerically weaker effects than in the above analyses.

### Supporting Tables and Figures

| Model | Range | $\alpha$ | $\beta$ | $\tau_{ALT}$ | $\tau_W$ | $e$ | $\omega$ |
|---|---|---|---|---|---|---|---|
| Model-free | 25th percentile | 0.80 | 5.06 | 0.53 | 0.17 | 0.71 | - |
| | Median | 1.00 | 9.59 | 0.90 | 0.62 | 0.83 | - |
| | 75th percentile | 1.00 | 15.47 | 1.00 | 1.00 | 0.90 | - |
| Model-based | 25th percentile | 0.55 | 5.36 | 0.69 | 0.82 | - | - |
| | Median | 0.97 | 8.08 | 0.83 | 0.88 | - | - |
| | 75th percentile | 1.00 | 14.84 | 0.99 | 0.96 | - | - |
| Hybrid | 25th percentile | 0.74 | 6.36 | 0.51 | 0.77 | 0.01 | 0.50 |
| | Median | 1.00 | 11.07 | 0.82 | 0.87 | 0.35 | 0.87 |
| | 75th percentile | 1.00 | 22.00 | 1.00 | 0.92 | 0.66 | 1.00 |

**Table S1. Reinforcement learning model parameter values.** Across-participant median and 25$^{th}$ and 75$^{th}$ percentile fit parameter values for the three models. $\alpha$ = learning rate; $\beta$ = softmax inverse temperature; $\tau_{ALT}$ = (1–decay) of non-chosen path values; $\tau_W$ = (1–decay) of non-presented world values; $e$ = eligibility trace (model-free and hybrid models only); $\omega$ = model-based weighting (hybrid model only).

| Model | -Lik | AIC | BIC |
|---|---|---|---|
| Model-free | 1948.43 | 4132.55 | 4488.93 |
| Model-based | 1624.75 | **3441.51** | **3726.61** |
| Hybrid | **1576.43** | **3441.44** | 3869.10 |

**Table S2. Reinforcement learning model fits.** Lower values indicate better fits to the behavioral data. The model-based model was an equivalent or better fit to the behavioral data than the hybrid model when accounting for additional parameters (as shown in the AIC and BIC values). For interpretation of the AIC measures, we consider the fit of the hybrid and model-based models to be equivalent, as the hybrid model was only 0.0017% better (using the model-free model as a baseline). -LIK = negative log likelihood (not penalized for number of parameters). AIC = Akaike Information Criterion; BIC = Bayesian Information Criterion.

| **Planning Period lag** | **Different start** | **State value** | - | - | Rarity |
| --- | --- | --- | --- | --- | --- |
| **Forward Current 70ms** | **(+) *** ** | **(+) ** ** | - | - | eq. |
| Forward Other 70ms | eq. | eq. | - | - | eq. |
| *Backward Current 40ms* | *eq.* | *eq.* | - | - | *eq.* |
| *Backward Other 40ms* | *eq.* | *eq.* | - | - | *eq.* |
| Forward Current 190ms | (+) p = 0.206 | eq. | - | - | eq. |
| Forward Other 190ms | (-) p = 0.144 | eq. | - | - | eq. |

| **Feedback Period lag** | Different start | - | Reward | RPE | **Rarity** |
| --- | --- | --- | --- | --- | --- |
| Forward Current 70ms | eq. | - | (-) p = 0.143 | eq. | eq. |
| Forward Other 70ms | eq. | - | eq. | eq. | eq. |
| Backward Current 40ms | (-) p=0.119 | - | eq. | eq. | eq. |
| **Backward Other 40ms** | eq. | - | eq. | eq. | **(+) *** |

**Table S3.** Full sequenceness regression results, including control analyses, in the planning period (top) and feedback period (bottom), for different state-to-state sequenceness lags. In each period, the replay direction and lag of interest is indicated in bold. In the planning period, we did not observe significant overall replay at a 40ms lag, but we use italics to report these control analyses for completeness. Effects equivalent to a null effect are indicated with 'eq.' (TOST procedure to rule out the presence of a medium- or larger-sized effect). The symbols (+) and (-) indicate the numerical direction of a relationship that is not equivalent to a null effect; p-values are directly noted or indicated with '*'. * p < 0.05; ** p < 0.01; *** p < 0.0001. We also conducted additional control analyses of the feedback period using data from the first 2.5 s of the period (excluding the first 160 ms to allow for initial processing). In this early feedback period, all relationships were equivalent to null effects.

| Planning | Region | T-statistic | Volume (mm3) | Peak Coord. |
|---|---|---|---|---|
| | – | – | 178,800 | – |
| | R Occipital lobe / Lingual gyrus | 7.07 | | 20, -81, -2 |
| | L Occipital lobe / Lingual gyrus | 5.76 | | -10, -61, -2 |
| | L Fusiform gyrus | 5.48 | | -30, -46, -17 |
| | R Thalamus / Pallidus | 4.96 | | 5, -1, 8 |
| | R Caudate | 4.06 | | 10, 4, 13 |
| | R Hippocampus | 3.92 | | 35, -26, -7 |
| | R Hippocampus / Anterior MTL | 2.88 | 300 | 30, -11, -22 |
| | **Hippocampus ROI** | | | |
| | R Hippocampus | 3.24 | 900 | 30, -36, -4 |
| **Current world** | R Hippocampus | 4.01 | 1125 | 25, -36, 3 |
| **Feedback** | **Region** | **T-statistic** | **Volume** | **Peak Coord.** |
| | – | – | 195,825 | – |
| | L Occipital lobe / Lingual gyrus | 6.99 | | -10, -91, -7 |
| | R Occipital lobe / Lingual gyrus | 6.86 | | 15, -86, -17 |
| | R Fusiform gyrus | 6.37 | | 30, -71, -17 |
| | L Fusiform gyrus | 6.27 | | -20, '-81, '-17 |
| | R Anterior Insula | 5.75 | | 30, 24, -2 |
| | R Caudate / Putamen | 5.1 | | 15, 9, 3 |
| | **Hippocampus ROI** | | | |
| | R Hippocampus | 3.63 | 825 | 30, -21, -12 |
| | L Hippocampus | 3.03 | 450 | -25, -16, -17 |
| | L Hippocampus | 3.02 | 225 | -15, -36, -2 |
| **Other world** | R Hippocampus | 3.23 | 75 | 25, -36, 3 |
| **Planning current > Feedback other** | | **T-statistic** | **Volume** | **Peak Coord.** |
| | R Fusiform gyrus | 3.32 | 2025 | 35, -51, 3 |
| | R Middle occipital gyrus | 3.32 | 375 | 25, -71, 3 |
| **Feedback other > Planning current** | | **T-statistic** | **Volume** | **Peak Coord.** |
| | R Inferior frontal gyrus / Insula | 3.47 | 2,475 | 25, 24, -2 |
| | L Medial frontal gyrus / OFC | 2.91 | 150 | -10, 14, -17 |
| | L Putamen | 2.97 | 150 | -25, 4, 3 |

**Table S4. Beamforming significant clusters.** Clusters significant at $p < 0.05$ using non-parametric permutation tests at the whole-brain level or within bilateral hippocampus ROI when noted. While we find larger areas of activity in the right hippocampus, it is possible that underlying bilateral activity is interpreted as unilateral activity by source reconstruction algorithms (O'Neill et al., 2021). For unthresholded maps, see https://neurovault.org/collections/11163/

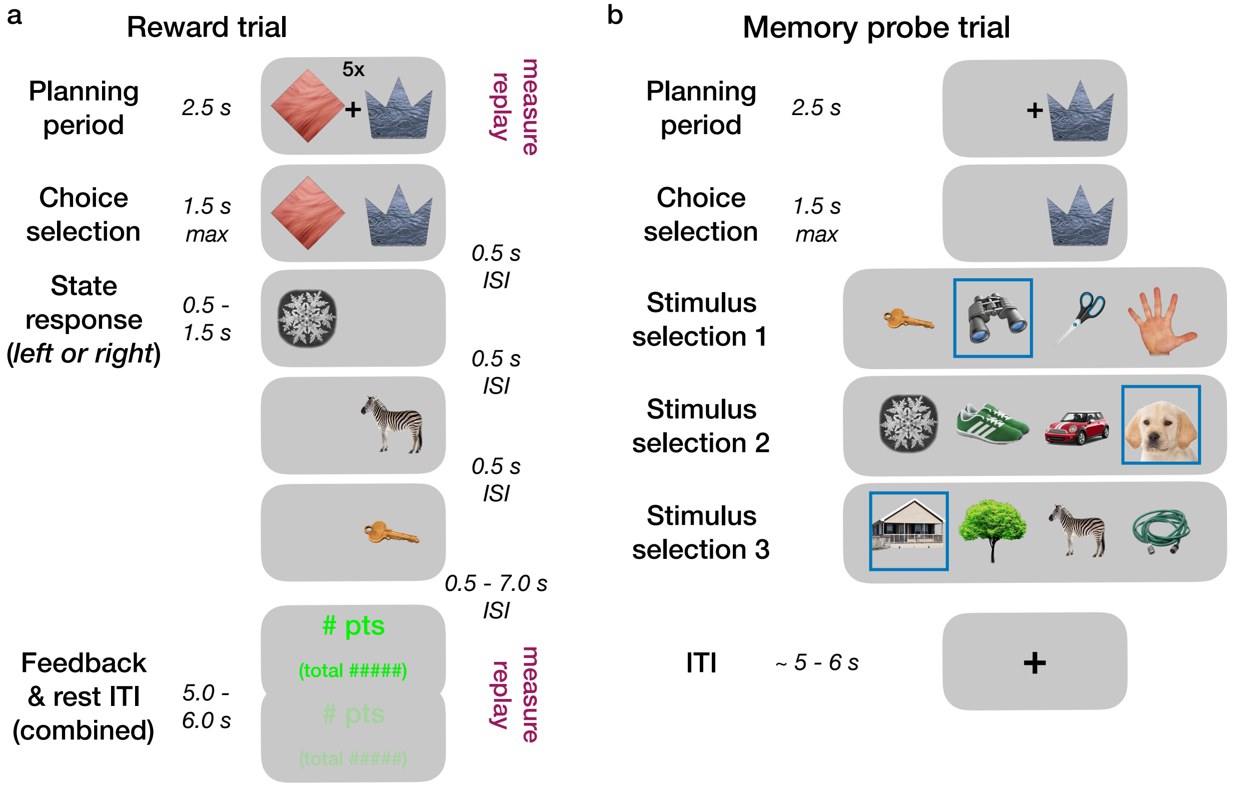


**Figure S1.** Trial structure and timing of reward learning trials and memory probe trials. (**a**) Trial timing of reward learning phase trials. Each trial began with a 2.5 s planning period, which was the focus on decision-related sequenceness analyses. Participants also saw the cued reward stakes above the shapes ('5x' or '1x'; see the Methods for a note on difficulty in interpreting effects of stakes in our data). Next, a central '+' disappeared and participants could enter their choice selection. Trials ended with a mean 5 s feedback and ITI period (with the total points, multiplied by the stakes, shown below). This period was the focus of feedback-related sequenceness analyses. There was no separate ITI in the scanned reward learning phase; the feedback faded out across time and served as an ITI. For the preceding non-rewarded structure learning phase where participants freely chose in order to learn the links between stimuli, the trial events and timing were as above, with the exception that no reward feedback was presented. Preceding the reward phase, structure learning phase trials (not shown) did not have reward feedback or per-trial stakes information. (**b**) Trial timing of memory test probe trials used in mini-blocks after structure learning and reward learning blocks. Participants were shown a single shape, and after selection of this shape, their goal was to select the correct next stimulus from four alternatives at each of the three stages.

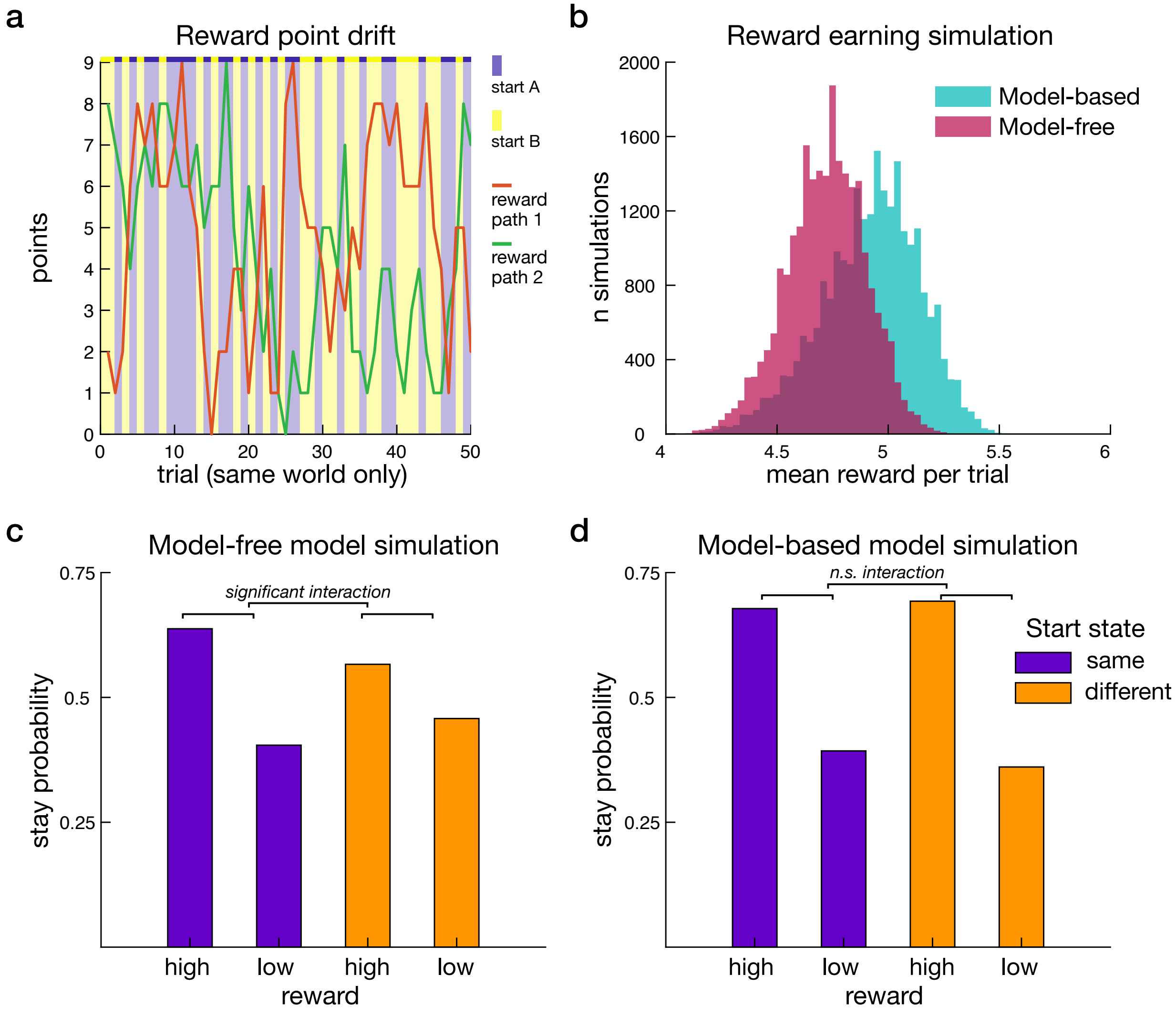


**Figure S2. Simulations of model-free and model-based RL models demonstrating effect of start state change on stay behavior and benefit of model-based behavior for reward earnings.** (**a**) Example reward point drift for the two options within one world, illustrating how a model-based agent can perform better than a model-free agent. The points associated with the two paths are shown in the red and green lines. The blue and yellow overlay represents the two alternative start states. Trials are concatenated to only show a single world; in the actual task, alternative world trials would be interspersed. (**b**) Simulations demonstrating the benefit of model-based behavior on mean reward earnings. Task parameters and reward distributions were as experienced by participants. 1000 simulations were conducted with each of the 24 individual participant's fit RL model parameters. Mean per-trial reward earnings were higher in the model-based model (4.919 [4.916 4.922]) than the model-free model (4.719 [4.716 4.721]). A given simulated model-based model earned more reward than a model-free model 79.9% of the time. (**c-d**) Simulation results of a group of 24 participants simulated 1000 times. Figures follow the format of behavioral results displayed in **Fig. 1d**. (**c**) Depiction of model-free model simulation results. In the same multilevel regression analysis as described in the main text, the interaction of reward by start state, indicating a model-free component to behavior, was significant in all simulations. (**d**). Depiction of model-based model simulation results. The interaction of reward by start state, indicating model-free component to behavior, was never significantly positive. Further, the interaction coefficient for every model-based group simulation was lower than all interaction coefficients across all model-free simulations. Both model-free and model-based simulations use the individual participant fit model parameters for each type of model and the underlying reward point drifts from the behavioral task, while regressions use the full graded reward feedback distribution. As for **Fig. 1d**, for display, reward points were binned into high and low reward, excluding the point value nearest the mean for this illustration. (Note that stay probabilities in a given condition will not add to 1 given that trials with near-mean feedback are excluded; this does not qualitatively affect the results.) As expected, the critical difference arises in the tendency to stay after reward when facing a different start state (orange bars) versus the same start state (purple bars). In the model-based model, the difference in stay probability is independent of start state, demonstrating generalization across specific top-level stimuli. The adaptive tendency to stay after reward versus stay after non-reward feedback when facing the same start state (purple bars) is weaker in the model-free model. Critically, the effect when the start state changed (orange bars) was much weaker in the model-free model. Note that in the model-free model, the residual effect of reward on stay choices when the start state changes (orange bars) is due to temporal autocorrelation in the drifting rewards associated with each path. I.e. a high reward on trial t when the start state changed is likely to have followed a high reward on trial t-1 when the start state did not change, allowing the model-free model to provide some appearance of adaptation when generalization is more beneficial.

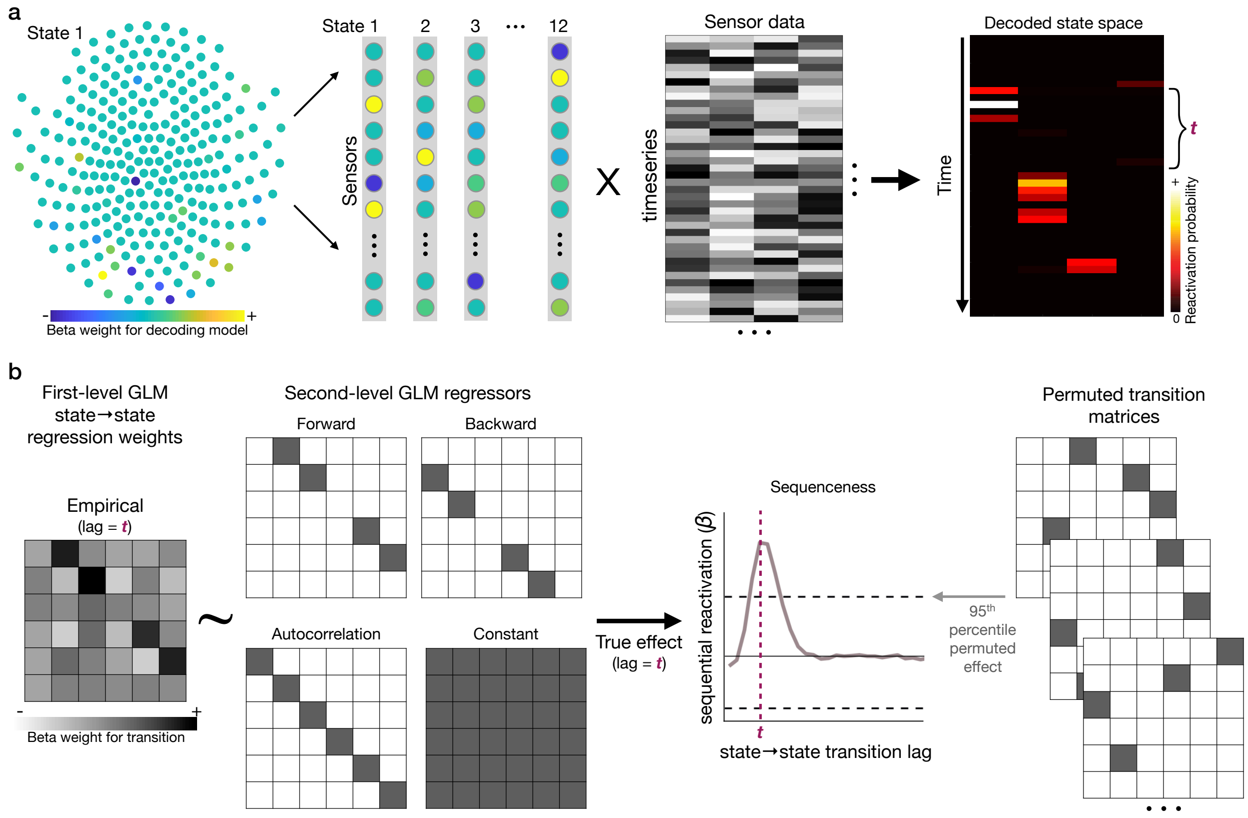


**Figure S3**. Schematic of sequenceness analysis pipeline. a) We train separate multivariate classifiers for each path state in the task from the localizer data, producing vectors of sensor weights for each state. The peak-accuracy classifiers are applied to the preprocessed MEG data [time x sensors] for each period in each trial (middle right). This transforms the data from sensor space to vectors of state reactivation [time x state]. For illustrative purposes only, we provide a visual example of sequential reactivation in the resulting state reactivation data. b) Using the state reactivation evidence data, a first-level GLM estimates the empirical strength of state-to-state transitions at each lag t for each period in each trial (left). For ease of visualization, only 6 states of a total 12 are depicted in the example matrices. This empirical matrix is the dependent variable in a second-level GLM, where the true forward and backward transition matrices are independent variables, alongside control autocorrelation and constant matrices (middle left). Per-trial results are z-scored across lags. For lag localization, results are then averaged across trials and across participants (middle right). To control for multiple comparisons, a non-parametric permutation approach is employed (right). The true state-to-state transition matrices are replaced by random transition matrices. To correct for multiple comparisons, a 95% threshold is applied to the maximum values across lags. Figure adapted from Liu et al. (Liu et al., 2021a) and Nour, Liu et al. (Nour et al., 2021).

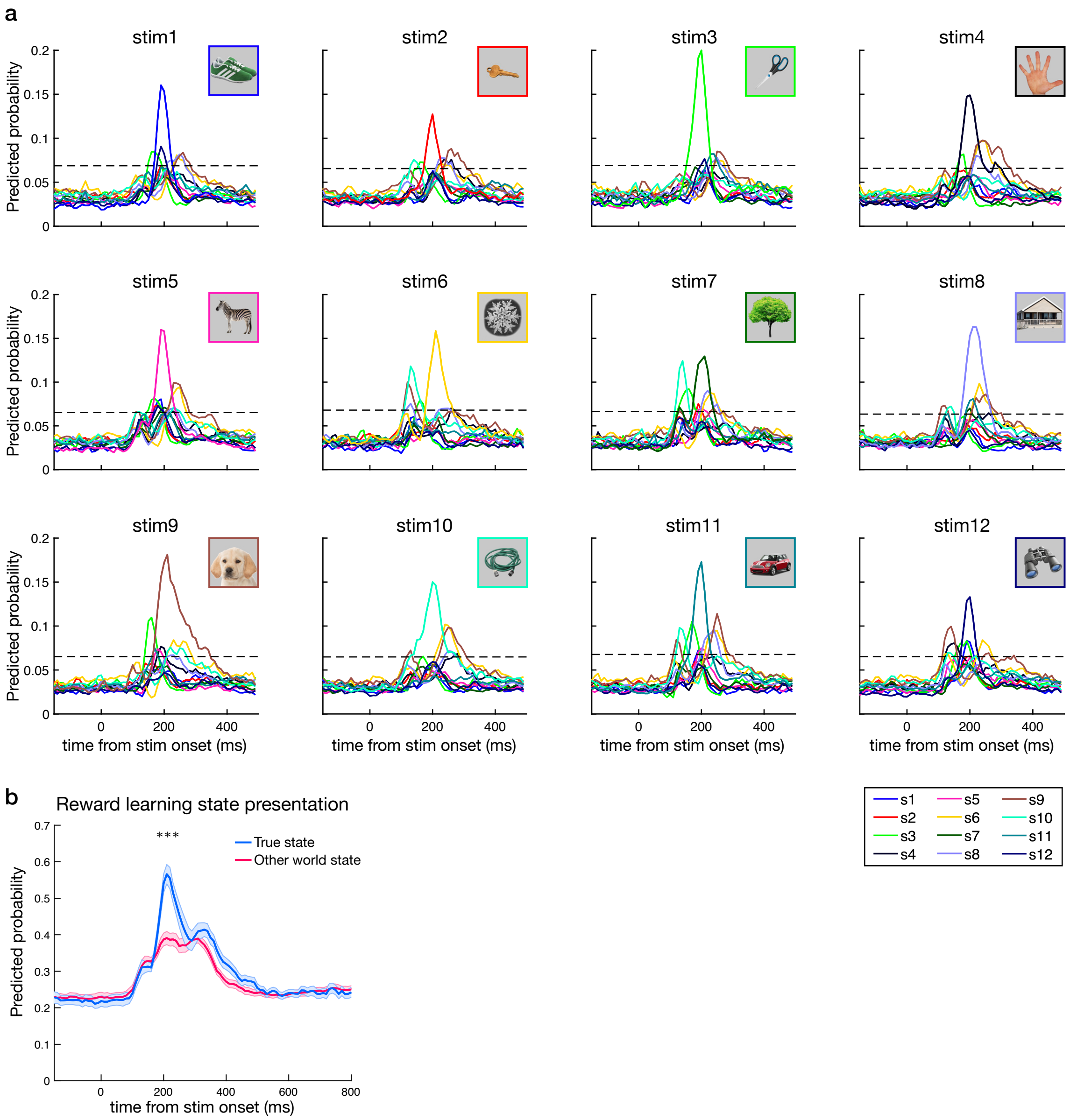


**Figure S4. Decoding performance of all 12 states during localizer and cross-classification of stimuli during reward learning.** (**a**) Cross-validated classification performance for each of the 12 path state stimuli in the localizer phase. Results represent training on the 200 ms time point and testing across all time points. Dotted lines indicate the permutation threshold estimated by randomly shuffling the labels and re-estimating the decoding. (Note that the actual stimulus 9 was a photo of the face of a young girl; the puppy stands in to avoid including identifiable information.) (**b**) Check of classifier performance when applied to actual stimuli during reward learning in the trial period where stimuli on each path are sequentially presented. Comparison of true versus other world stimuli centered around 200 ms training time of classifier (190-210 ms; *** $p < 10^{-7}$ t-test true state versus non-presented state; peak at 220 ms). Shaded error margins represent SEM.

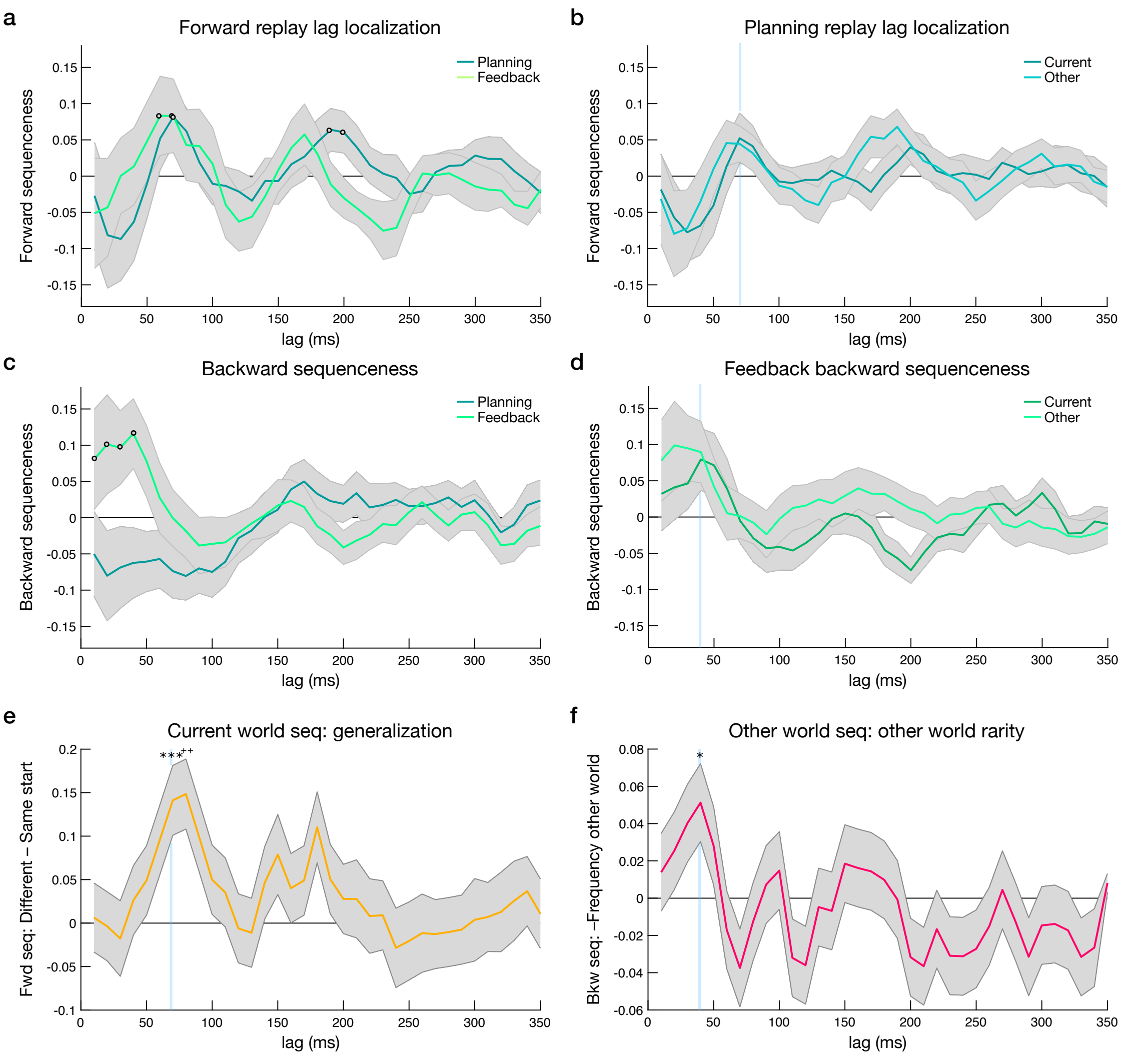


**Figure S5. Sequenceness and regressions for the full range of time lags.** (**a**) Forward sequenceness for all learned paths during planning and feedback periods was evident for a common state-to-state lag of 70 ms in both trial periods, and 190-200 ms during planning alone. Open dots indicate time points exceeding a permutation significance threshold, which differs for the two periods. (**b**) Planning period forward sequenceness separately for current world and other world paths. (**c**) Backward sequenceness for all learned paths during planning and feedback periods was evident for state-to-state lags that spanned 10-50 ms during the feedback period alone. Open dots indicate time points exceeding a permutation significance threshold, which differs for the two periods. (**d**) Feedback period forward sequenceness separately for current and other world paths. (**e**) Planning forward sequenceness and generalization: no alternative lag exceeded that of the effect at 70ms, and no alternative lag effect was significant after correcting for multiple comparisons (at a lag of 180ms, the effect exceeded an uncorrected threshold). (**f**) Feedback period replay and rarity: no alternative lag exceeded that of the effect at 40ms, and no alternatives were significant at an uncorrected or corrected level. For all panels, note that the x-axis in the sequenceness panels indicates the lag between reactivations, derived as a summary measure across seconds; the axis does not represent time within a trial period.

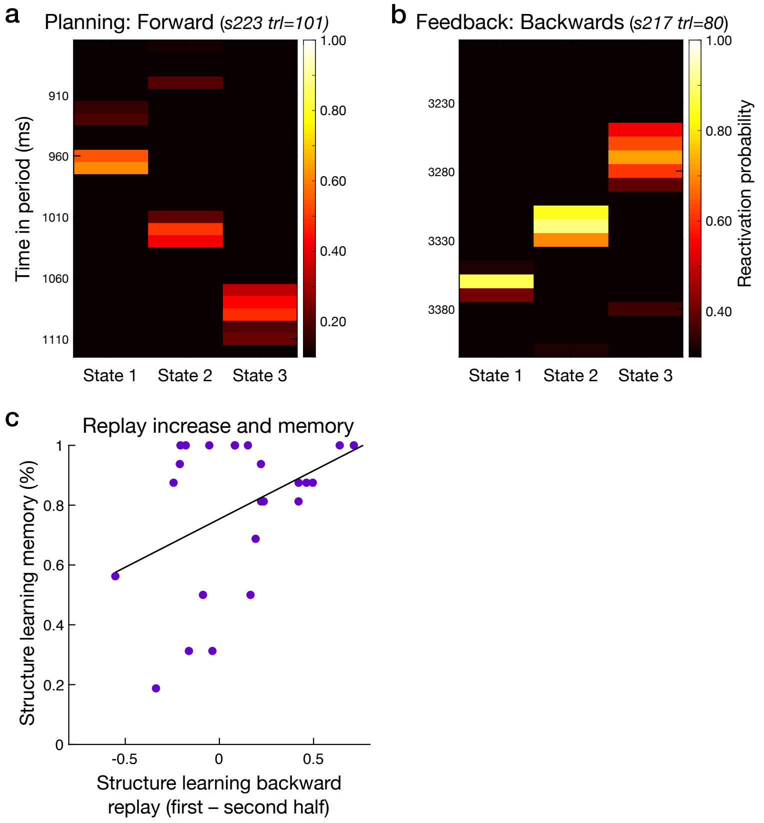


**Figure S6. Individual sequenceness example events and individual difference correlations.** (**a**) Example of forward sequenceness with a 70 ms state-to-state lag in the planning period for current world paths. (**b**) Example of backward sequenceness with a 40 ms state-to-state lag in the feedback period for other world paths; s = participant; trl = trial. Time is relative to the onset of planning or feedback. Note that these are presented as examples only; the sequenceness linear regression method, which provides the sequenceness measures used in all our analyses, utilizes data across the full time period of interest and does not identify individual events. (**c**) Structure learning phase: relationship between increased backward replay during the inter-trial interval and memory performance in the structure learning phase (p = 0.047).

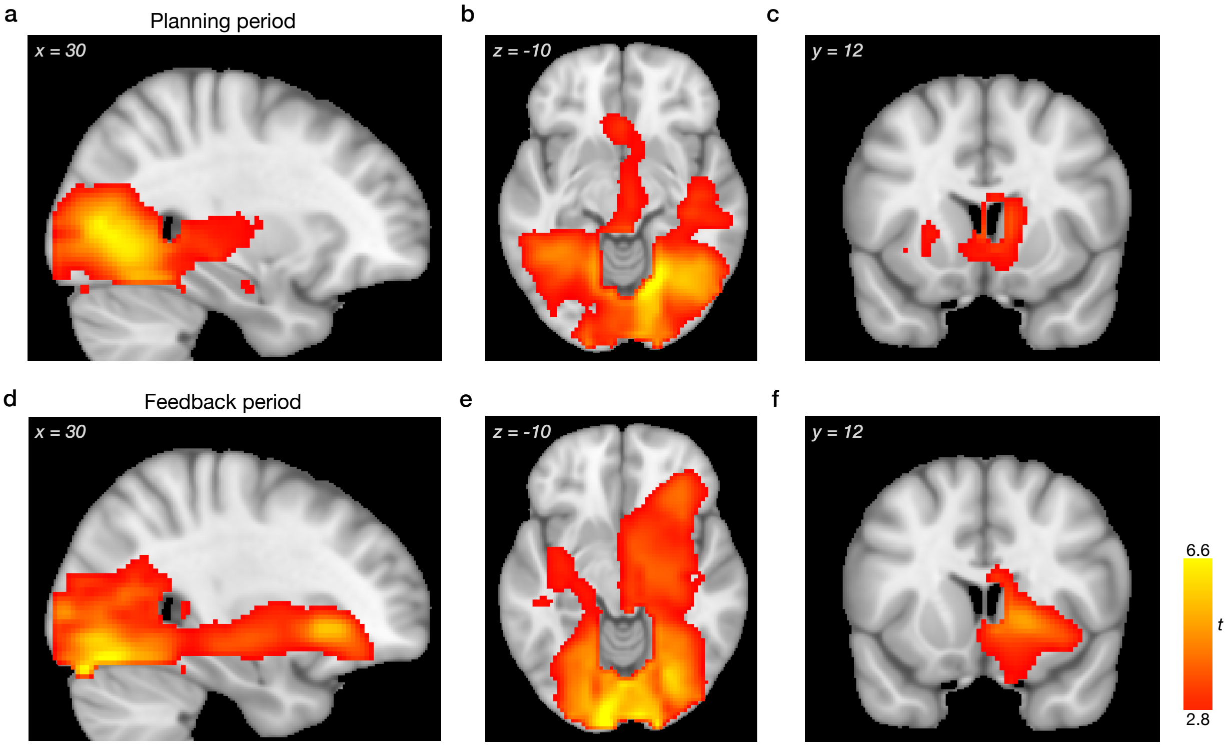


**Figure S7. Additional views of replay onset source localization (beamforming).** (**a-c**) Power increases associated with replay onset in the planning period in the MTL (a-b) and striatum (c). (**d-f**) Power increases associated with replay onset in the reward feedback period in the MTL (d-e) and striatum (f). For display, statistical maps were thresholded at $p < 0.01$ uncorrected; clusters significant at $p < 0.05$, whole-brain corrected using non-parametric permutation test. For unthresholded statistical maps and results within the hippocampus ROI mask see https://neurovault.org/collections/11163/.

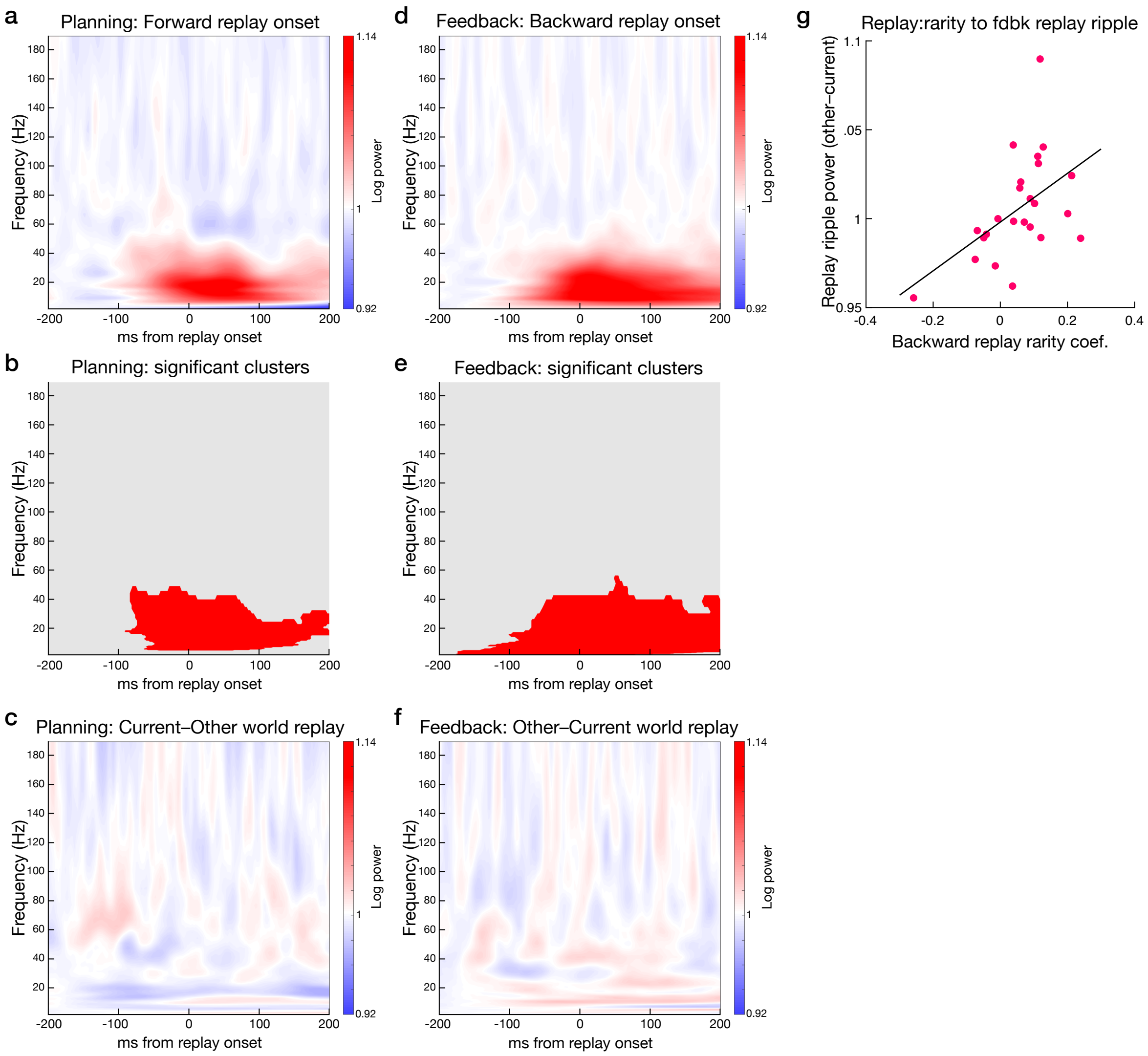


**Figure S8. Replay onset full-frequency results.** (**a**-**c**) Planning period. (**a**) Power increases associated with replay onset in the planning period across the full 0-190 Hz range. (**b**) Significant power increases in the planning period in red. Clusters significant at $p < 0.05$; corrected using non-parametric 2D permutation test, after an initial thresholding at $p < 0.01$. (**c**) Planning replay onset power difference for current world paths minus other world paths; no significant differences. (**d**-**f**) Feedback period. (**d**) Power increases associated with replay onset in the reward feedback period across the full 0-190 Hz range, as in panel a. (**e**) Significant power increases in the reward feedback period, as in panel b. (**f**) Feedback replay onset power difference for other world paths minus current world paths; no significant differences. (**g**) Feedback period ripple frequency band ROI for other minus current world paths correlated with individual differences in the feedback replay memory rarity effect. However, overall, we saw no main effect of replay-associated power increase at higher frequencies (Liu et al., 2019; Liu et al., 2021b; Nour et al., 2021), with changes more consistent with previous lower-frequency power increases observed for replay of episodic memories (Wimmer et al., 2020).

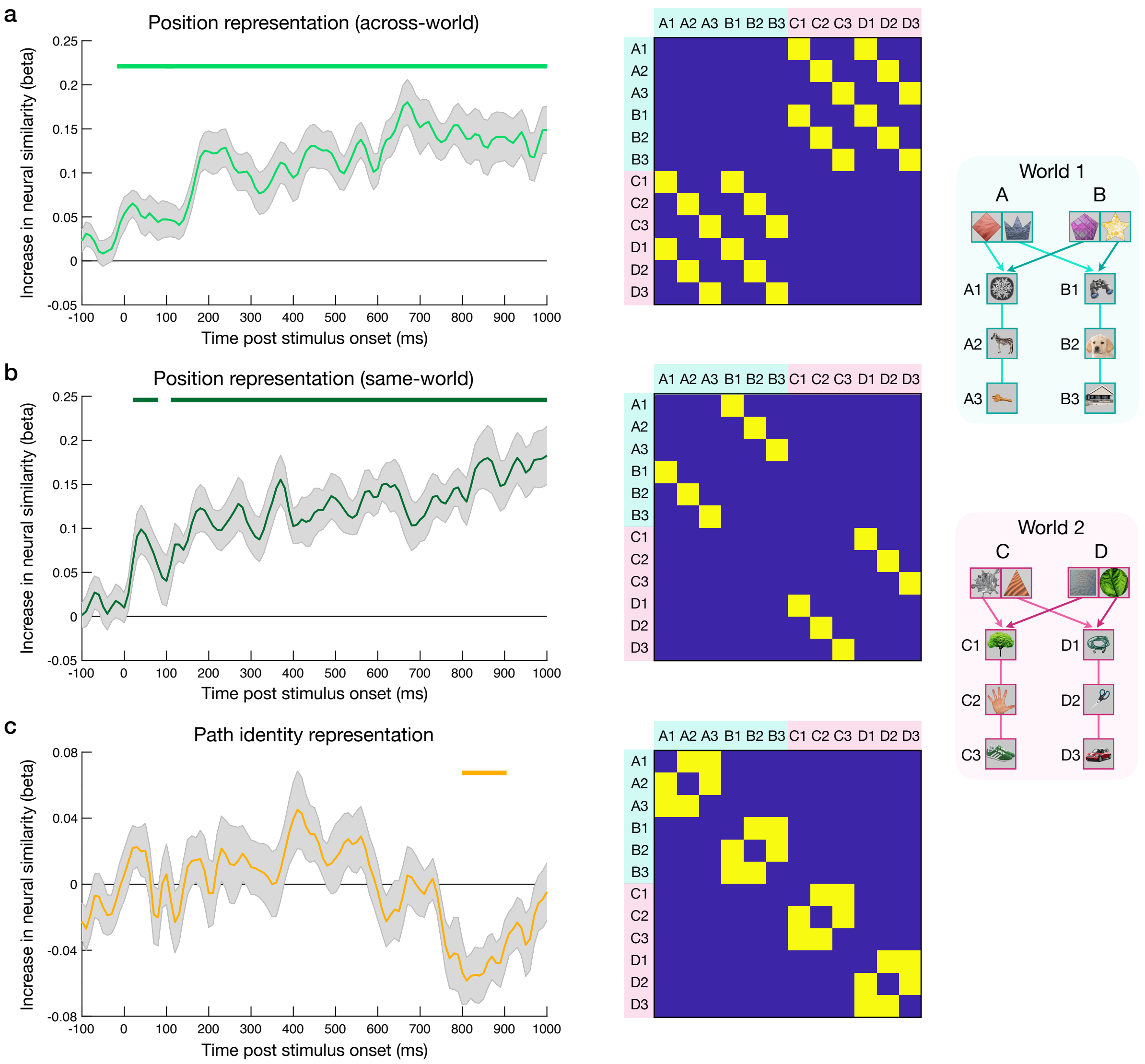


**Figure S9. Change in neural similarity after learning.** Labeled task diagram, right column of figure. (**a**) Stimuli occupying the same path position increased their similarity to stimuli at the same position in the alternative world, left. Across-world position design matrix, middle. (**b**) Stimuli at the same path position increased their similarity to stimuli at the same position in the same world, left. Same-world position regression matrix, right. (**c**) Path identity: stimuli within the same path decreased their similarity, left. Path identity regression matrix, right. Note that position representations do not return to baseline because learning task trials continue on to other fixed-order stimuli. Specifically, in visual presentation, state1 is followed by state 2 and 3, and thus a position 1 representation transitions into a representation for position 2, while a position 2 representation transitions into position 3.

**SI References**


Barr, D.J., Levy, R., Scheepers, C., and Tily, H.J. (2013). Random effects structure for confirmatory hypothesis testing: Keep it maximal. *J Mem Lang 68*.

Carey, A.A., Tanaka, Y., and van der Meer, M.A.A. (2019). Reward revaluation biases hippocampal replay content away from the preferred outcome. *Nat Neurosci 22*, 1450-1459.

Collins, A.G., and Frank, M.J. (2012). How much of reinforcement learning is working memory, not reinforcement learning? A behavioral, computational, and neurogenetic analysis. *Eur J Neurosci 35*, 1024-1035.

Daw, N.D., Gershman, S.J., Seymour, B., Dayan, P., and Dolan, R.J. (2011). Model-based influences on humans' choices and striatal prediction errors. *Neuron 69*, 1204-1215.

Daw, N.D., O'Doherty, J.P., Dayan, P., Seymour, B., and Dolan, R.J. (2006). Cortical substrates for exploratory decisions in humans. *Nature 441*, 876-879.

Deuker, L., Bellmund, J.L., Navarro Schroder, T., and Doeller, C.F. (2016). An event map of memory space in the hippocampus. *Elife 5*.

Diedrichsen, J., and Kriegeskorte, N. (2017). Representational models: A common framework for understanding encoding, pattern-component, and representational-similarity analysis. *PLoS Comput Biol 13*, e1005508.

Doll, B.B., Duncan, K.D., Simon, D.A., Shohamy, D., and Daw, N.D. (2015). Model-based choices involve prospective neural activity. *Nat Neurosci 18*, 767-772.

Eldar, E., Roth, C., Dayan, P., and Dolan, R.J. (2018). Decodability of reward learning signals predicts mood fluctuations. *Current biology : CB*.

Gupta, A.S., van der Meer, M.A., Touretzky, D.S., and Redish, A.D. (2010). Hippocampal replay is not a simple function of experience. *Neuron 65*, 695-705.

Kool, W., Cushman, F.A., and Gershman, S.J. (2016). When does model-based control pay off? *PLoS Comput Biol 12*, e1005090.

Kool, W., Gershman, S.J., and Cushman, F.A. (2017). Cost-benefit arbitration between multiple reinforcement-learning systems. *Psychol Sci 28*, 1321-1333.

Kurth-Nelson, Z., Economides, M., Dolan, R.J., and Dayan, P. (2016). Fast sequences of non-spatial state representations in humans. *Neuron 91*, 194-204.

Lakens, D. (2017). Equivalence tests: A practical primer for t tests, correlations, and meta-analyses. *Soc Psychol Personal Sci 8*, 355-362.

Lau, B., and Glimcher, P.W. (2005). Dynamic response-by-response models of matching behavior in rhesus monkeys. *J Exp Anal Behav 84*, 555-579.

Liu, Y., Dolan, R.J., Higgins, C., Penagos, H., Woolrich, M.W., Olafsdottir, H.F., Barry, C., Kurth-Nelson, Z., and Behrens, T.E. (2021a). Temporally delayed linear modelling (tdlm) measures replay in both animals and humans. *Elife 10*.

Liu, Y., Dolan, R.J., Kurth-Nelson, Z., and Behrens, T. (2019). Human replay spontaneously reorganises experience. *Cell 178*, 640-652 e614.

Liu, Y., Mattar, M.G., Behrens, T.E.J., Daw, N.D., and Dolan, R.J. (2021b). Experience replay is associated with efficient nonlocal learning. *Science 372*.

Luyckx, F., Nili, H., Spitzer, B., and Summerfield, C. (2019). Neural structure mapping in human probabilistic reward learning. *Elife 8*.

Nour, M.M., Liu, Y., Arumuham, A., Kurth-Nelson, Z., and Dolan, R.J. (2021). Impaired neural replay of inferred relationships in schizophrenia. *Cell*.

O'Neill, G.C., Barry, D.N., Tierney, T.M., Mellor, S., Maguire, E.A., and Barnes, G.R. (2021). Testing covariance models for meg source reconstruction of hippocampal activity. *Sci Rep 11*, 17615.

Patzelt, E.H., Kool, W., Millner, A.J., and Gershman, S.J. (2019). Incentives boost model-based control across a range of severity on several psychiatric constructs. *Biol Psychiatry 85*, 425-433.

Schuirmann, D.J. (1987). A comparison of the two one-sided tests procedure and the power approach for assessing the equivalence of average bioavailability. *J Pharmacokinet Biopharm 15*, 657-680.

Sutton, R.S., and Barto, A.G. (1998). Reinforcement learning: An introduction (Cambridge: MIT Press).

Van Veen, B.D., Van Drongelen, W., Yuchtman, M., and Suzuki, A. (1997). Localization of brain electrical activity via linearly constrained minimum variance spatial filtering. *IEEE Transactions on biomedical engineering 44*, 867-880.

Wikenheiser, A.M., and Redish, A.D. (2015). Hippocampal theta sequences reflect current goals. *Nat Neurosci 18*, 289-294.

Wimmer, G.E., Braun, E.K., Daw, N.D., and Shohamy, D. (2014). Episodic memory encoding interferes with reward learning and decreases striatal prediction errors. *J Neurosci 34*, 14901-14912.

Wimmer, G.E., Li, J.K., Gorgolewski, K.J., and Poldrack, R.A. (2018). Reward learning over weeks versus minutes increases the neural representation of value in the human brain. *J Neurosci 38*, 7649-7666.

Wimmer, G.E., Liu, Y., Vehar, N., Behrens, T.J., and Dolan, R.D. (2020). Episodic memory retrieval success is related to rapid replay of episode content. *Nat Neurosci*.

Wu, C.T., Haggerty, D., Kemere, C., and Ji, D. (2017). Hippocampal awake replay in fear memory retrieval. *Nat Neurosci 20*, 571-580.
